# Loss of consciousness reduces the stability of brain hubs and the heterogeneity of brain dynamics

**DOI:** 10.1101/2020.11.20.391482

**Authors:** Ane López-González, Rajanikant Panda, Adrián Ponce-Alvarez, Gorka Zamora-López, Anira Escrichs, Charlotte Martial, Aurore Thibaut, Olivia Gosseries, Morten L. Kringelbach, Jitka Annen, Steven Laureys, Gustavo Deco

## Abstract

Low-level states of consciousness are characterised by disruptions of brain dynamics that sustain arousal and awareness. Yet, how structural, dynamical, local and network brain properties interplay in the different levels of consciousness is unknown. Here, we studied the fMRI brain dynamics from patients that suffered brain injuries leading to a disorder of consciousness and from subjects undergoing propofol-induced anaesthesia. We showed that pathological and pharmacological low-level states of consciousness displayed less recurrent, less diverse, less connected, and more segregated synchronization patterns than conscious states. We interpreted these effects using whole-brain models built on healthy and injured connectomes. We showed that altered dynamics arise from a global reduction of network interactions, together with more homogeneous and more structurally constrained local dynamics. These effects were accentuated using injured connectomes. Notably, these changes lead the hub regions to lose their stability during low-level states of consciousness, thus attenuating the core-periphery structure of brain dynamics.

## 1 Introduction

It is widely accepted that consciousness is decreased during sleep, under anaesthesia, or as a consequence of major brain lesions producing disorders of consciousness (DOC). In clinical settings, different states of consciousness have been defined depending on the level of wakefulness and awareness [1], as measured by the responsiveness and the ability to interact with the environment. The study of these different levels of consciousness has proved to be essential to understand the neural correlates of consciousness, yet, the underlying mechanisms remain largely unknown. Elucidating these mechanisms is challenging since they seemingly rely on a non-trivial combination of alterations in local dynamics and network interactions.

During the last decades, the study of the organization of brain dynamics and connectome structure has provided increased understanding of the healthy brain structure and function [2, 3, 4, 5, 6, 7]. On the one hand, analyses of electroencephalography (EEG), functional MRI (fMRI), and magnetoencephalography (MEG) have shown that a hallmark of healthy awake brain dynamics is the balance between integration and segregation [8, 9, 10, 11]. On the other hand, graph theory studies have shown that the modular and hierarchical organization of the human connectome optimizes the efficiency and robustness of information transmission [3, 12]. For these reasons, consciousness has been considered to result from the interplay between dynamics and connectivity allowing the coordination of brain-wide activity to ensure the conscious functioning of the brain [13, 14, 15, 16]. In contrast, unconscious states are characterized by a loss of integration [17, 14, 18], a loss of functional complexity [19, 20], and a loss of communication at the whole-brain level [21, 22, 9, 18]. Interestingly, it has been shown that the repertoire of functional correlations is more constrained by the anatomical connectivity during unconscious states [23, 24, 25, 13]. In other words, the dependency of dynamics on structural connections is increased in low-level states of consciousness. Along with these network effects, it has been proposed that some local brain regions, such as fronto-parietal regions, posterior cingulate, precuneus, thalamus and parahippocampus, play an important role in maintaining consciousness [1, 26, 27]. To study how structural, dynamical, local and network brain properties interplay in the different levels of consciousness, theoretical models are needed to incorporate all these levels of description.

In this study, we built whole-brain models with global and local parameters to investigate the possible mechanisms underlying the reduction of consciousness as a consequence of severe brain injury and transient physiological modifications due to anaesthesia. For this, we studied the fMRI dynamics of patients who have suffered brain injuries from various etiologies (i.e. traumatic brain injury (TBI), anoxia, haemorrhage) affecting different brain regions implicated in DOC. Specifically, we analysed data from patients with Unresponsiveness Wakefulness Syndrome (UWS; preserved arousal but no behavioural signs of consciousness)[28] and in Minimally Conscious State (MCS, fluctuating but reproducible signs of consciousness) [29], and compared them with healthy control subjects (CNT) during wakefulness. We also considered the fMRI dynamics of healthy controls scanned during conscious wakefulness (W), during propofol-induced anaesthesia (state of deep sedation, S) and during the recovery from it (R). To study the brain dynamics, we used phase-synchronization analyses, which have proven to effectively describe the spatiotemporal dynamics of fMRI signals [30]. We interpreted the results using a whole-brain model based on Hopf bifurcations [31]. This model is able to generate different collective oscillatory dynamics depending on the (healthy or injured) anatomical connectivity structure, the global strength of connections and the local state of the network’s nodes. Importantly, the model allows the investigation of the interplay between structure and global and local dynamics. In particular, it allows to relate the network behaviors to the local dynamics of regions having an important topological role in the network, such as the structurally highly connected nodes, or “hubs”.

## 2 Results

We performed both data- and model-driven analyses to compare different levels of consciousness in two neuroimaging datasets comprising DOC and healthy subjects under anaesthesia. The first dataset consisted of fMRI signals and structural connectomes (SC) from healthy subjects during conscious wakefulness (n =35), and MCS and UWS patients (n = 33 and n = 15, respectively). The analysis was complemented with an additional fMRI data from 16 healthy controls scanned during conscious wakefulness (W), deep sedation (S) and recovery from it (R).

### Decrease in brain data-driven phase dynamics complexity in low-level states of consciousness

We first searched for spatiotemporal signatures of loss of consciousness in the whole-brain blood-oxygen-level-dependent (BOLD) signals for the different experimental groups. We were interested in synchronization dynamics, thus, we concentrated on phase statistics. For this, following previous research, BOLD phases were extracted in the 0.04-0.07 Hz frequency band [32, 6, 30] using the Hilbert transform (Fig. 1 a-b). This allows to obtain, at each time point *t*, a phase-interaction matrix, *P*(*t*), given by the phase differences among the regions of interest (ROIs) (Fig. 1 c, see Methods). We were interested in the spatiotemporal organization of these phase interactions.

**Figure 1:**
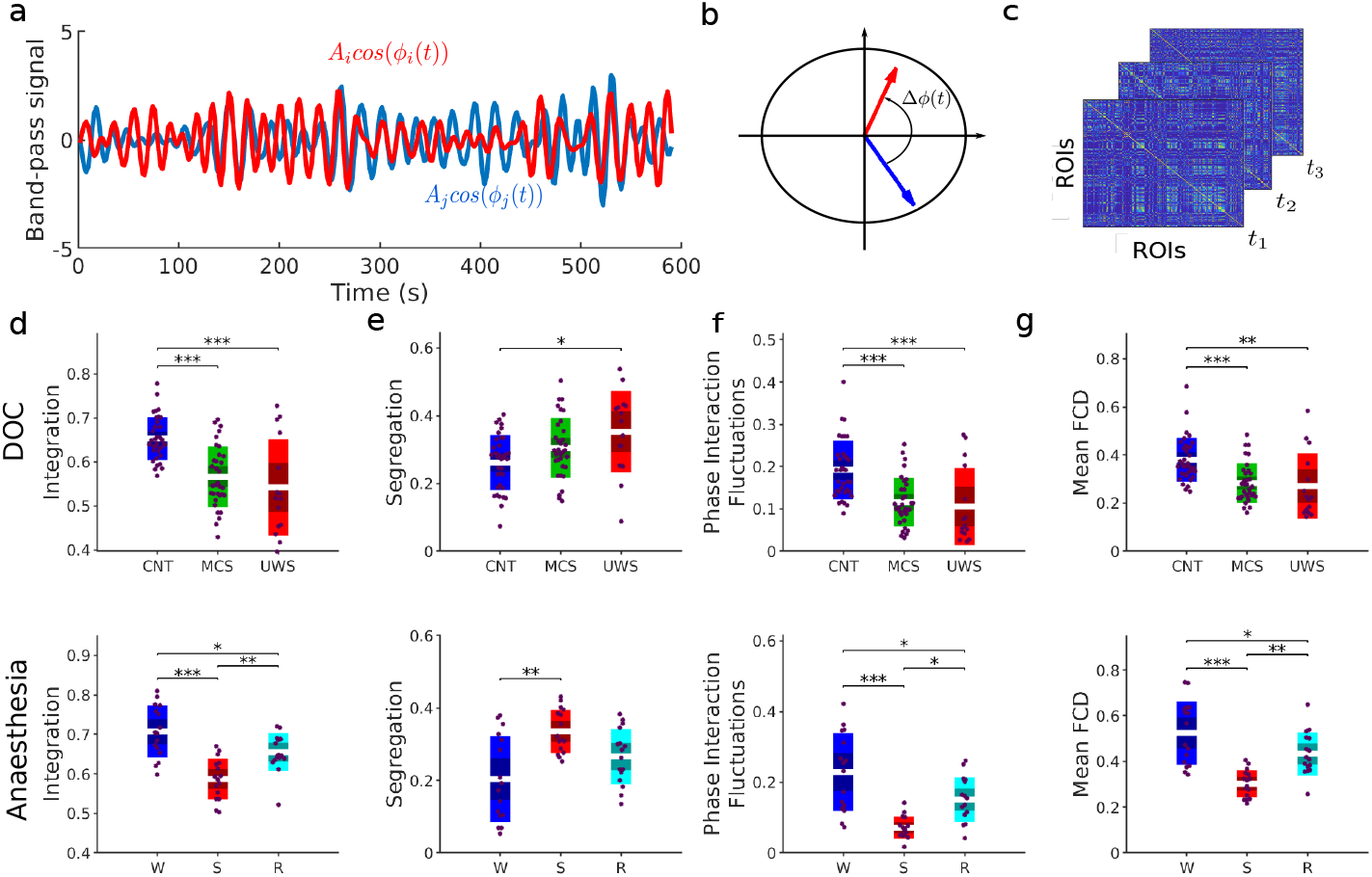
Changes in global properties of phase-dynamics induced by loss of consciousness. **a)** BOLD band-pass signals (0.04-0.07 Hz) for two samples ROIs. The instantaneous phases, *φ_j_* (*t*) and *φ_k_* (*t*), of each signal were computed using the Hilbert transform. **b)** At each time frame, the interaction between ROIs was given by the instantaneous phase difference, *Δφ_jk_* (*t*) = |*φ_j_* (*t*) — *φ_k_* (*t*)|, which can be represented as vectors in the unit circle of the complex plane. **c)** Phase-interaction matrices *P_jk_*(*t*) were calculated as the cosine of the phase difference, cos(Δ*φ_jk_*(*t*)), at time t. All global measures used afterwards were based on the phase-interaction matrices. **d-e)** The structure of phase interactions was described in terms of the integration and the segregation of the time-averaged phase interaction matrix (see Methods). **f)** We quantified the temporal fluctuations of the mean phase synchrony (i.e., the average over ROIs of matrix *P*(*t*)) through its temporal standard deviation. **g)** To detect the existence of recurrent synchronization patterns, we computed the FCD comparing phase-interaction matrices at different time (see Methods). Briefly, the FCD represents the (cosine) similarities between phase-interaction matrices at times *t* and *t*’ for all possible pairs (*t*, *t*’). The panel shows the average similarity for each experimental condition. In panels d-g, each dot represents a participant and the boxes represent the measure’s distribution. Differences between groups were assessed using one-way ANOVA followed by FDR p-value correction. *: *p* < 0.05; **: *p* < 0.01; ***: *p* < 0.001 (see Table 1 for details).

First, we examined the spatial distribution of phase-interaction matrices. We measured the integration and segregation of the phase-interaction matrices averaged over time. Integration was measured as the size of the largest subcomponent. Segregation was measured as the modularity index of the matrix resulting from community detection (see Methods). We found that the average integration across time was significantly lower for MCS and UWS, compared to CNT, and for S and R compared to W (Fig 1 d, see also table 1). For the average segregation, we observed the opposite pattern (Fig 1 e, see also table 1). Thus, low-level states of consciousness were characterised by a decrease of integration and an increase of segregation.

**Table 1:** Results of the mean values of the global measurements for each group and statistics. Statistics were computed with a one-way-ANOVA, followed by FDR correction (adjusted p-values are shown). The table shows the mean values and standard error of the empirical measures of integration, segregation, phase interaction fluctuations and mean FCD.

| DOC Datasets | Integration | Segregation | Phase Interaction Fluctuations | Mean FCD |
| --- | --- | --- | --- | --- |
| CNT | 0.653 ± 0.008 | 0.26 ± 0.01 | 0.19 ± 0.01 | 0.38 ± 0.02 |
| MCS | 0.56 ± 0.01 | 0.31 ± 0.01 | 0.12 ± 0.01 | 0.28 ± 0.02 |
| UWS | 0.54 ± 0.03 | 0.35 ± 0.03 | 0.11 ± 0.02 | 0.27 ± 0.03 |
| ANOVA | $p < 0.001$<br>$F_{2,80} = 18.51$ | $p = 0.006$<br>$F_{2,80} = 5.21$ | $p < 0.001$<br>$F_{2,80} = 13.39$ | $p < 0.001$<br>$F_{2,80} = 10.9$ |
| Multiple Comparisons |  |  |  |  |
| $p_{CNT-MCS}$ | < 0.001 | 0.2072 | < 0.001 | 0.014 |
| $p_{CNT-UWS}$ | < 0.001 | 0.016 | < 0.001 | 0.023 |
| $p_{MCS-UWS}$ | 0.241 | 0.223 | 0.882 | 0.594 |
| Anaesthesia Datasets | Integration | Segregation | Phase Interaction Fluctuations | Mean FCD |
| W | 0.71 ± 0.02 | 0.20 ± 0.03 | 0.23 ± 0.03 | 0.52 ± 0.04 |
| S | 0.59 ± 0.01 | 0.34 ± 0.02 | 0.07 ± 0.01 | 0.30 ± 0.01 |
| R | 0.65 ± 0.01 | 0.27 ± 0.02 | 0.15 ± 0.02 | 0.43 ± 0.02 |
| ANOVA | $p < 0.001$<br>$F_{2,45} = 18.8$ | $p < 0.001$<br>$F_{2,45} = 8.93$ | $p < 0.001$<br>$F_{2,45} = 17.46$ | $p < 0.001$<br>$F_{2,45} = 18.8$ |
| Multiple Comparisons |  |  |  |  |
| $p_{W-S}$ | < 0.001 | 0.001 | < 0.001 | < 0.001 |
| $p_{W-R}$ | 0.029 | 0.126 | 0.014 | 0.041 |
| $p_{S-R}$ | 0.006 | 0.115 | 0.014 | 0.004 |

Second, we evaluated the temporal fluctuations of the average phase-interaction matrix. For this, we computed the standard deviation of the mean phase-interaction value across time, providing an estimate of the diversity of the average synchronization (see Methods). We found significant reduction of phase-interaction fluctuations in low-level states of consciousness compared to conscious states (Fig. 1 f, top; see also table 1).

Temporal fluctuations of the average phase-interaction matrix indicate excursions of the total level of synchronization over time but, alone, they do not capture the presence of recurrent connectivity patterns. Therefore, we next evaluated the temporal recurrence of phase-interaction matrices over time, or functional connectivity dynamics (FCD, see Methods), that describes how recurrent in time the synchronization patterns were. Briefly, this method computes the phase-interaction matrices averaged in sliding time window of 30 s and measures the similarity across all pairs of time windows, which is summarized in the FCD matrix. We found that low-level states of consciousness presented a significantly lower mean FCD value than in normal wakefulness (Fig. 1 g; see also table 1 and Supplementary Fig. 1). This suggests that phase configurations were less recurrent in time for low-level states of consciousness.

Altogether, the above results show that, in both pathological and pharmacological low-level states of consciousness, brain phase-synchronization patterns were less connected, less diverse and less recurrent in time than in healthy conscious states.

### Decreased model-based global functional connectivity in low-levels states of consciousness

To gain insights into the possible mechanisms underlying the above changes in BOLD phase statistics, we studied a whole-brain computational model. Because we were interested in synchronous oscillations, we modelled the local dynamics of single brain regions using the normal form of a Hopf bifurcation (see Methods, Eqs. 5 and 6). By varying a single bifurcation parameter *a_j_*, local dynamics of a brain region *j* can transit from noisy oscillations (*a_j_* < 0) to sustained oscillations (*a_j_* > 0) (Supplementary Fig. 2). The frequency of oscillations was estimated from the peak of the BOLD power spectral density in the frequency band 0.04-0.07 Hz. The dynamics of the *N* = 214 brain regions were coupled through the connectivity matrix *C_jk_*, which was given by the connectome of healthy subjects. The matrix *C_jk_* was scaled by the global coupling *g*. Thus, the large-scale network was weakly or strongly connected for small or large values of *g*, respectively (Fig. 2 a). In summary, at this level of description the network dynamics depended on three ingredients: the local parameters for each node (*a_j_*), the global strength of connections (*g*) and the network’s structure (*C_jk_*).

**Figure 2:**
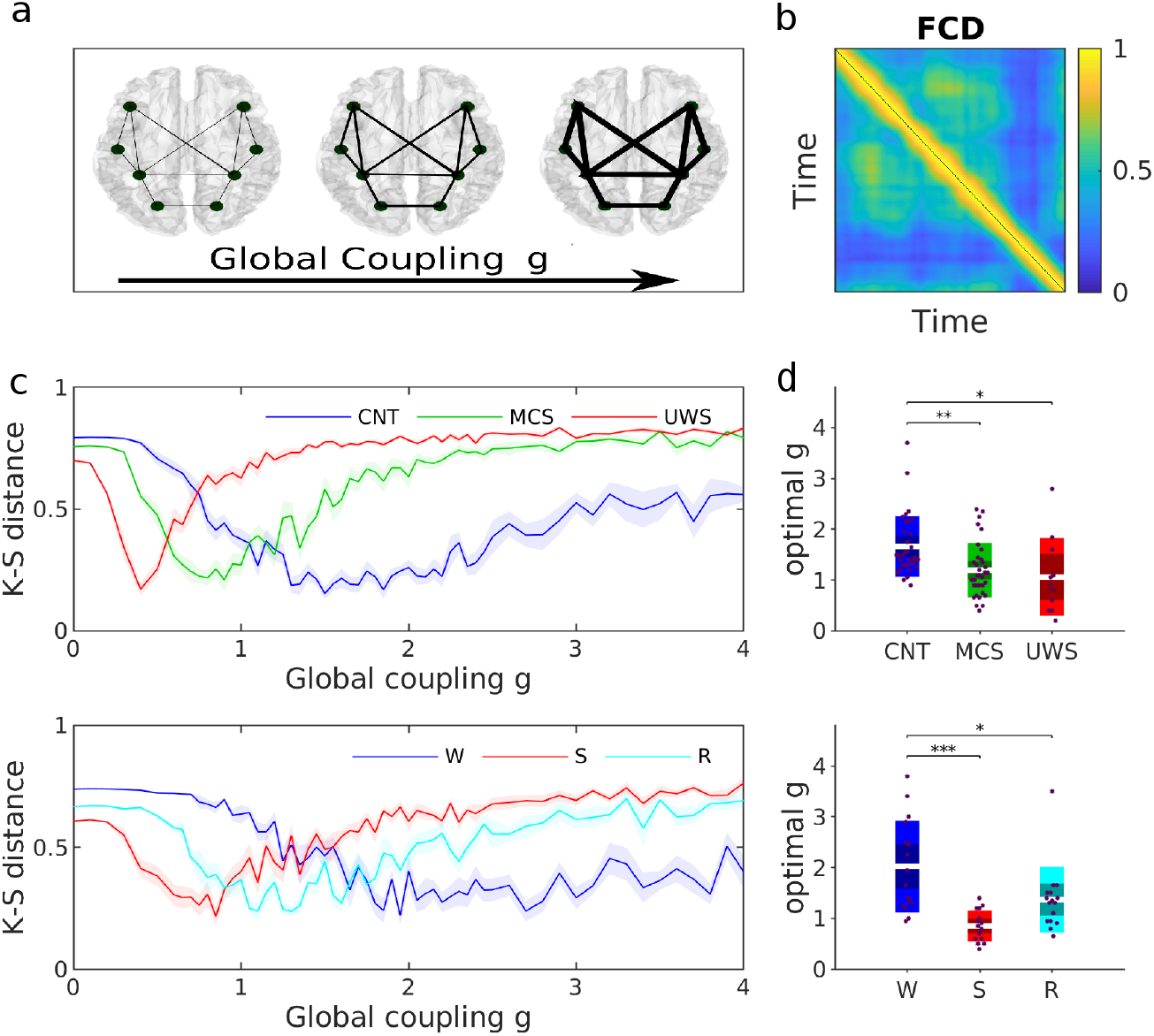
Fitting of global coupling parameter in the whole-brain network model. **a)** The global coupling model parameter *g* scales the weights of the SC matrix. Low and high values of *g* represent weakly and strongly coupled networks, respectively. **b)** To estimate this global parameter, we sought for the model that best reproduced the distribution of FCD values (fixing all other model parameters). **c)** KS-distance between the empirical and the model FCD distributions, as a function of *g*, for one participant of each subject group (top: healthy controls and DOC patients; bottom: awake and anaesthetised subjects). Solid lines and shaded areas represent the mean and the standard error of the fitting curves over simulation trials. **d)** Optimal global coupling *g* for all participants. In each panel, each dot represents a participant and the boxes represent the distribution of *g*. Differences between groups were assessed using one-way ANOVA followed by FDR p-value correction. *: *p* < 0.05; **: p < 0.01; ***: p < 0.001. In panels c and d, we used the healthy structural connectome as the underlying connectivity of all models.

First, we studied the network dynamics for the homogeneous case, in which we set *a_j_* = 0 for all nodes. This choice was based on previous studies which suggest that the best fit to the empirical data arises at the brink of the Hopf bifurcation where *a* ~ 0 [31]. In this case, the network dynamics were determined by a single free parameter, the global coupling strength *g*. This parameter was estimated by fitting the FCD statistics of the data, as in previous studies [31, 33]. Specifically, for each experimental condition, we evaluated the agreement between the simulated and the empirical group FCD using the Kolmogorov-Smirnov distance (KS-distance, Fig. 2 b). The KS-distance reached a minimum at different values of *g* for the different experimental conditions, with *g* being the lowest for states of low-levels of consciousness (Fig 2 c-d, see table 2). Notably, although the fit of the model was based on the FCD, the models also maximized the fit of other data statistics, such as the FC and the metastability (Supplementary Fig. 3).

**Table 2:** Estimated global couplings for all experimental conditions. p-values: *P_CNT-MCS_* = 0.015, *P_CNT-UWS_* = 0.019, *P_MCS-UWS_* = 0.7984; *P_W-S_* < 0.001, *P_W-R_* = 0.031, *P_R-S_* = 0.080.

| Conditions | CNT | MCS | UWS | W | S | R |
| --- | --- | --- | --- | --- | --- | --- |
| Global coupling $g$ | $1.7 \pm 0.1$ | $1.2 \pm 0.1$ | $0.8 \pm 0.2$ | $2.0 \pm 0.2$ | $0.9 \pm 0.1$ | $1.4 \pm 0.2$ |

Furthermore, we found that the increase in global coupling strength *g* with consciousness goes in line with a decrease in the correlation between the structural and functional connectivities (Supplementary Fig. 4). These results indicate that in low-level states of consciousness the brain dynamics were more constrained by the structural connectivity due to lower coupling values. Indeed, low global coupling restricts the network interactions to ROIs directly connected by a link, while increasing the global coupling favours the propagation of activity within the network, and gives rise to correlations between nodes indirectly coupled.

### Heterogeneous model

We next asked whether we can obtain additional information by relaxing the local bifurcation parameter which enforced all ROIs to operate at the same working point. We studied the heterogeneous case in which the local parameters *a_j_* were allowed to vary. The individual parameters *a_j_* were estimated from the data using a gradient descent method (see Methods). In this model, the *g* parameter was fixed to the one previously estimated with the homogeneous model (i.e., all *a_j_*=0). We note that the resulting distribution of local parameters contributed to network collective dynamics, since shuffling the values of *a* across brain regions lead to worse fits of the network statistics (Supplementary Fig. 5).

We inspected the estimated bifurcation parameters across nodes within and across groups. We found that bifurcation parameters in normal wakefulness (i.e., CNT and W) tended to be more negative compared to low-level states of consciousness (Fig. 3 a-d), i.e., they tended to display more stable noisy oscillations. Notably, this was specially the case for the structural hub ROIs, which showed strong negative values of the local bifurcation parameter *a* during normal wakefulness (Fig 3 a-d, Supplementary Fig. 6). When comparing normal wakefulness before anaesthesia (W) and recovery from anaesthesia (R) we found a similar distributions of the bifurcation parameter *a* values. In particular, the negativity was reestablished for hubs (Fig 3 c-d). This tendency was also observed when comparing MCS and UWS (Supplementary Fig. 7) and, even if in these cases the difference in the bifurcation parameter values was smaller than for the others, the hubs had more negative values in MCS. Using linear stability analysis, we showed that the hubs have a stabilizing role within the dynamical system, i.e., they contribute to the most stable eigenvectors, and lose their stability for low-level states of consciousness (see Supplementary Information and Supplementary Fig. 8). These results show that the hubs lost their stabilizing role in low-level states of consciousness.

We computed the difference in local parameters between patients and controls and between anaesthesia/recovery and wakefulness (Fig. 3 e). The highest absolute difference in local parameters between controls and MCS/UWS patients were found in subcortical regions, such as the thalamus, caudate, and hippocampus, the amygdala, and in cortical regions such as calcarine, insula, fusiform, frontal superior orbital, precuneus, cingulum, and temporal areas (Fig 3 f top left and Supplementary Tables 1-2). When comparing the local parameters for wakefulness and anaesthesia, the regions with the highest absolute difference values included subcortical regions such as the thalamus, caudate, hippocampus, parahippocampal, and putamen, and cortical regions such as cingulum, insula, and some regions of the frontal part, paracentral and precentral (Fig. 3 f top right and Supplementary Table 3). Finally, the highest differences between wakefulness and recovery were found in the hippocampus, the cingulum and the precuneus (Fig. 3 f bottom right and Supplementary Table 4).

**Figure 3:**
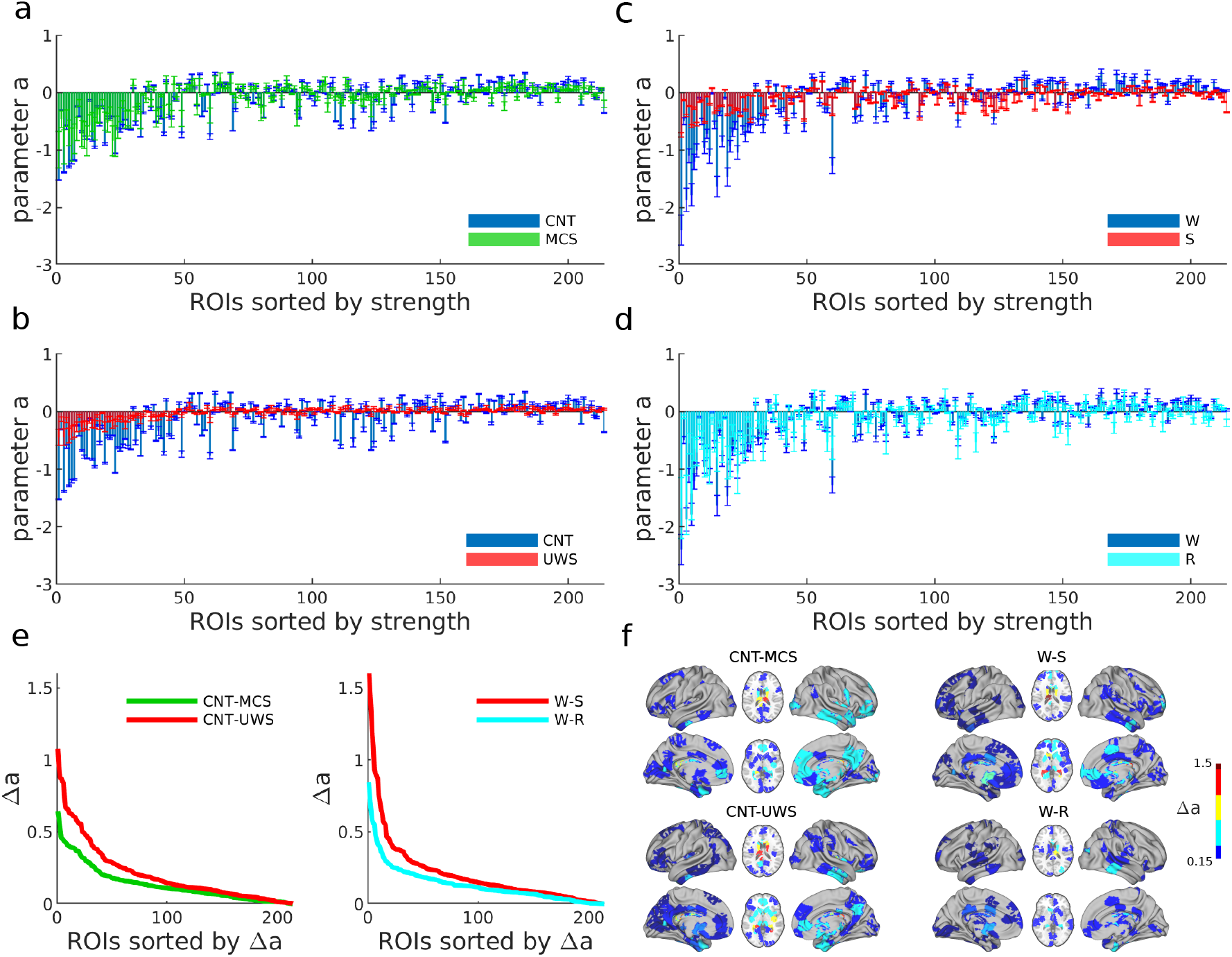
Local bifurcation parameters of the whole-brain model. **a-d)** Estimated bifurcation model parameters *a* for each of the 214 nodes (sorted by node strength). Bars indicate the mean ± standard deviations across simulation trials. Results for low-level states of consciousness (MCS and UWS) are compared against the healthy controls in a) and b). Results for anesthesia and recovery (S and R states) are compared to the initial awake state (W) in c) and d) respectively. **e)** Ranked absolute parameter difference, Δ*a*, for all the comparisons. **f)** Spatial distribution of Δ*a* > 0.15 in the brain for each of the group comparisons.

### Loss of heterogeneity in low-level states of consciousness

In general, if we observe the dynamics of a node within a network and we estimate its node-specific parameters, these parameters are affected by the network interactions, because we only have access to the dynamics of the ROIs *embedded* in the network. In the following, we used a strategy to disentangle the changes in local parameters due to network effects from those due to local modifications. This analysis provides information about the origin (local or network-related) of the different dynamics of the ROIs for the different states of consciousness.

Indeed, one can define an *effective* local parameter composed of the bifurcation parameter (*a_j_*) and the connectivity strength of each node (*S_j_* =Σ_*k*_ *C_jk_*), given as: 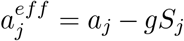, (see Methods). Thus, for the family of homogeneous models (*a_j_* = *const*.), the effective parameter is linearly related to the connectivity strength, while, in the heterogeneous case, we expect deviations from this linear relation. In other words, in the homogeneous case, differences in effective local dynamics are fully explained by the network connections. In contrast, the heterogeneous case can produce additional diversity of local dynamics.

We used this relation to distinguish between homogeneous and heterogeneous dynamics in the different data associated with different levels of consciousness. First, we estimated the effective bi-furcation parameters 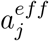 from the data in each brain state using gradient descent with fixed *g* for each condition (the values of *g* were those of Fig. 2 d). Note that, in this case, instead of estimating *a_j_*, the method estimates directly 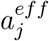 (see Methods, Eq. 10). Next, we evaluated the deviation from the linear relation between the estimated effective bifurcation parameter and the strength of the nodes (Fig. 4 a-b). We found that linear regression residuals were larger for control subjects and during healthy wakefulness than for DOC patients and anaesthesia (Fig. 4 c, *p* < 0.001 for all comparison in both dataset (computed with a one-way-ANOVA, followed by FDR correction, with the exception of *p_S-R_* = 0.002). This means that, on one hand, conscious states were associated with more heterogeneous dynamics for which different brain regions had different local dynamics. On the other hand, low-level states of consciousness were associated with homogeneous dynamics for which differences in local dynamics were explained to a large degree by differences in connectivity strength alone.

**Figure 4:**
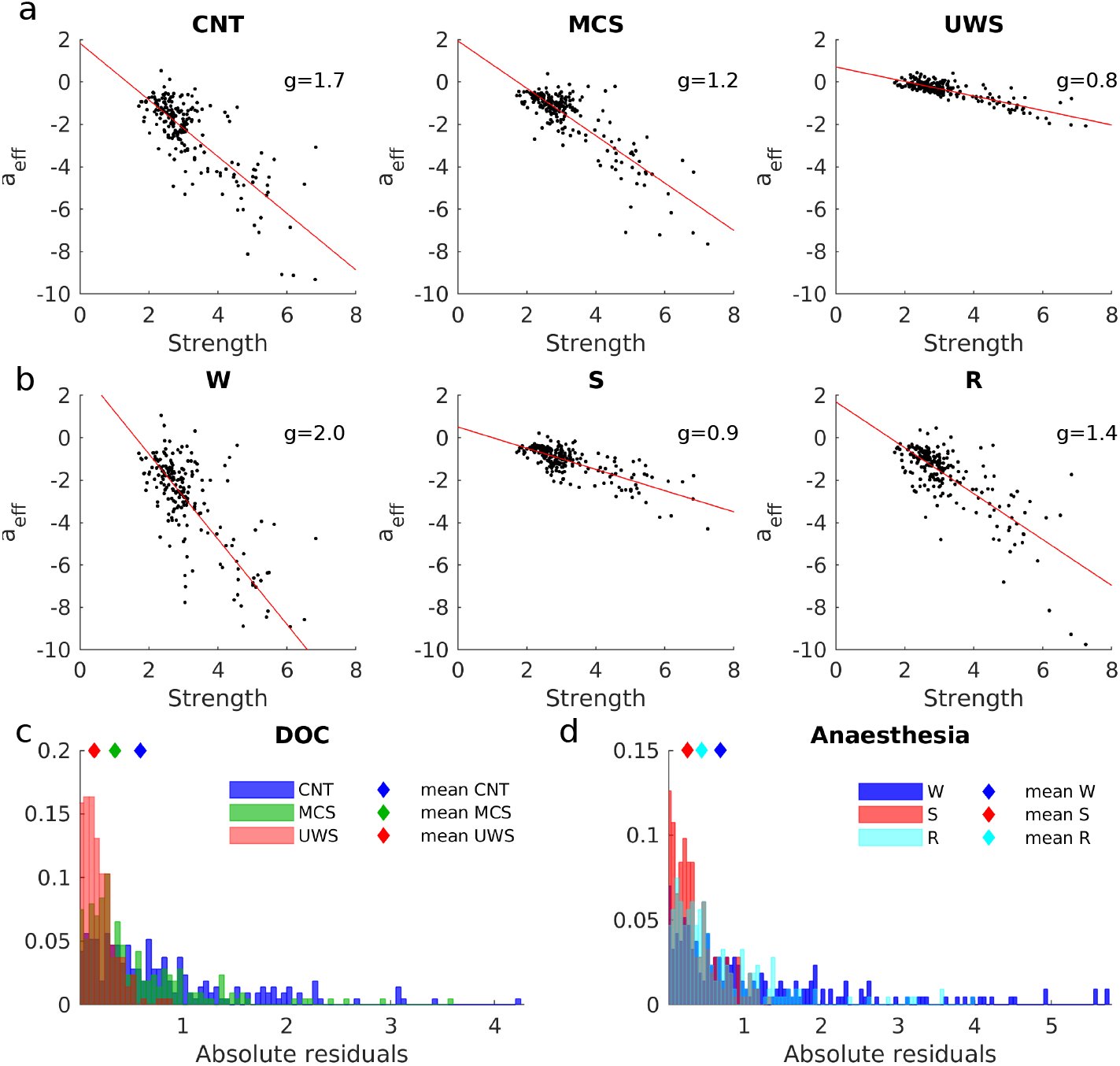
Disentangling structurally- and dynamically-driven heterogeneity of local nodes. **a-b)** The effective local bifurcation parameters, 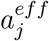, were estimated using the heterogeneous model. In this model, the parameters 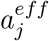 were optimized, after fixing *g* to that obtained for the homogeneous model (see Methods). The obtained parameters were compared to the strengths of the nodes *S_j_*, for healthy controls and DOC patients (a) and for awake and anesthetized conditions (b). In each panel, each dot represents one node. The red lines indicate the linear fits. **c)** Distribution of the absolute residuals of each node given by the squared difference between the value of 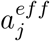 and the estimated linear relationship between 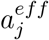 and *S_j_*, for each group. **d)** Same as c) but for W, S and R states.

### Alteration of the structural connectivity core in DOC patients

Up to now, the structural connectivity of the models was given by the connectome of healthy subjects. This allowed the study of dynamical factors leading to loss of consciousness. In the following, we studied the effect of injured anatomical connectivity on brain dynamics by considering the connectomes from DOC patients (Fig. 5 a and Supplementary Fig. 9). First, we quantified the alterations in connectomes caused by brain injuries through the strength of the nodes and the network’s rich-club architecture.

**Figure 5:**
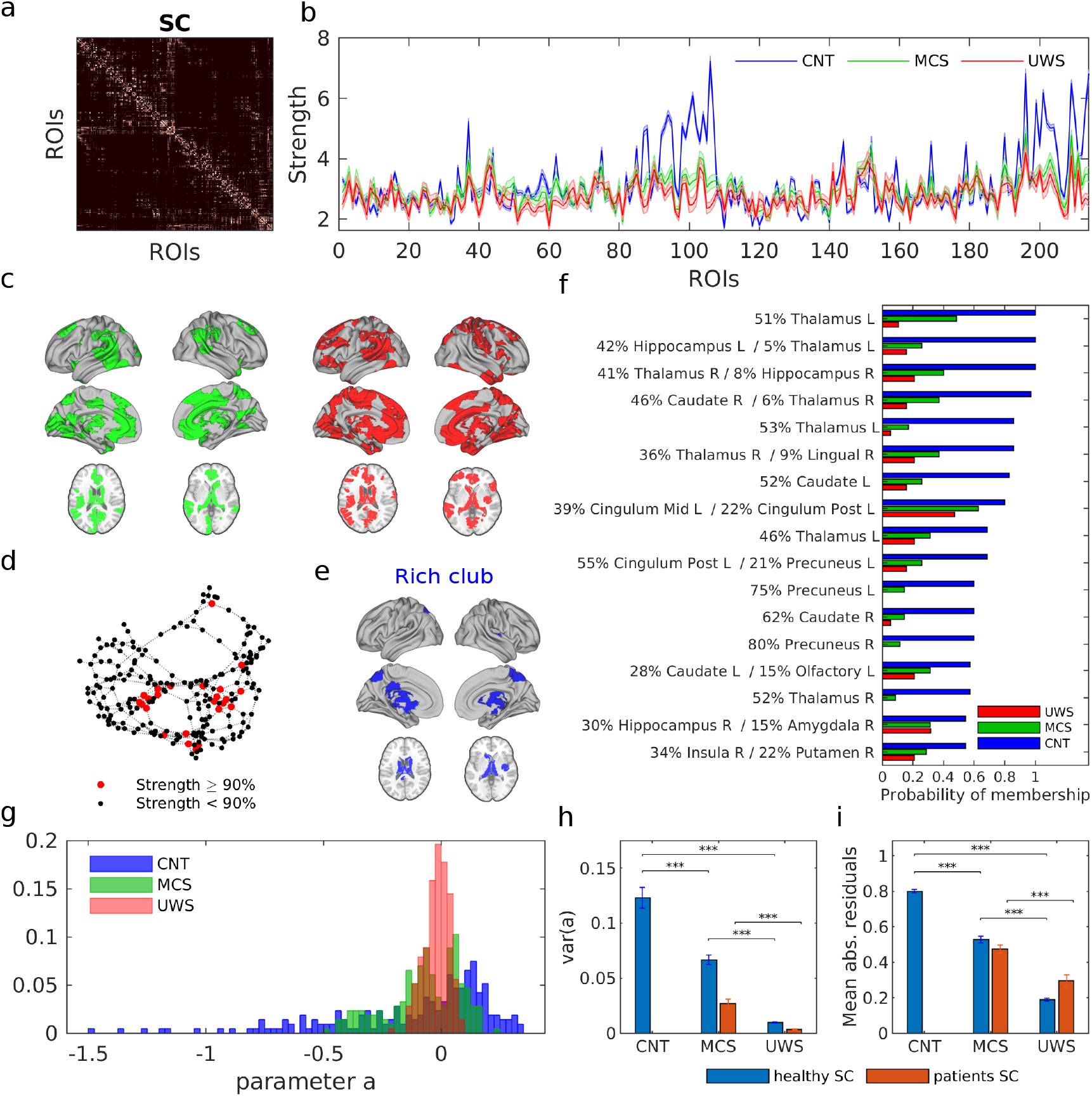
Disruption of the structural connectome in DOC patients. **a)** SC matrices were averaged over subjects for each clinical group (CNT, MCS, and UWS). **b)** Average node strength of each node for each group. Shaded areas represent the standard error across subjects. **c)** ROIs with significant differences in strength between controls and patients (Wilcoxon rank sum test, followed by FDR correction). Top (green): CNT-MCS comparison; bottom (red): CNT-UWS comparison. **d)** Visualization of hubs: each node corresponds to one ROI and the edges correspond to the SC (only connections >0.2 are shown). The graph was displayed using force-directed layout, i.e. attractive and repulsive forces between strongly and weakly connected nodes, respectively. Highly connected nodes, i.e. with high strength, are called structural hubs (red dots). **e)** Hubs which are highly connected among themselves form a rich club (RC) sub-network (here depicted in blue for the average SC of control subjects). **f)** For each ROI, the probability of forming part of the RC was computed by RC identification in the individual SC matrices (see Methods). The percentage corresponds to the covered part of the ROI in the AAL parcellation.**g)** Distribution of the estimated bifurcation parameters *a_j_* using the average SC for each clinical group (healthy, MCS, and UWS). **h)** The variance of the distribution of parameters *a_j_* for each clinical group. **i)** Median of the absolute residuals of the linear relationship between the *a_eff_* vs strength. ***: *p* <0.001, Wilcoxon rank sum test, followed by FDR correction.

We found that the strength of each ROI, i.e. the sum of connections of one node averaged over subjects, significantly decreased for DOC patients compared to controls for several brain regions (p < 0.05, Wilcoxon rank sum test with FDR correction; Fig. 5 b). These regions included the thalamus, the posterior and the anterior cingulum, hippocampus, the frontal medial, motor areas, caudate, precuneus, insula and precentral, for MCS patients (Fig. 5 c left, and see also Supplementary Table 5) and the aforementioned ones plus the fusiform, the parahippocampal, the cuneus, the lingual and the temporal areas for UWS patients (Fig 5 c right, and see details in Supplementary Table 6).

We next examined the interconnections between the ROIs with larger strength by the detection of a rich-club organization (see Methods). A network is said to contain a rich-club when (*i*) it contains hubs and (*ii*) those hubs are densely interconnected among themselves forming a cluster. (Fig. 5 d, see also Supplementary Fig. 10). In the healthy SC, the rich club was composed mostly of subcortical (thalamus, hippocampus, and caudate) and cortical regions such as the insula, the precuneus and the posterior cingulum (Fig. 5 e). We calculated the probability of each ROI to pertain to the rich club across individual subjects for controls and DOC patients (see Methods). We found that, when present in DOC patients, rich clubs were made of less ROIs and their composition varied from subject to subject (Fig. 5 f and Supplementary Fig. 11). Overall, these results show that connectomes from DOC patients presented alterations in the formation and the composition of the rich-clubs.

Finally, we studied the global and local dynamics of whole-brain models with large-scale connections constrained by the injured connectomes. Using these connectomes, we did not find significant differences in the global coupling parameter *g* (Supplementary Fig. 12). This was due to the high inter-individual variability of the structural connectomes. In contrast, consistent with the results above, we found that heterogeneity of local dynamics was reduced for models corresponding to DOC patients (Fig. 5 g-h and Supplementary Fig. 13). Moreover, the dynamically-based heterogeneity was significantly reduced (Fig. 5 i and Supplementary Fig. 14), indicating that local parameters were strongly determined by structural connections. These effects were stronger using injured SCs than using the healthy SC for all conditions, indicating that structural damage additionally impairs the emergence of heterogeneity.

## 3 Discussion

In the present study, we analysed and modelled brain dynamics from patients that show reduced consciousness due to brain damage (MCS and UWS), to propofol-induced anaesthesia (S), and to recovery from it (R). We showed that reduction of consciousness is characterized by brain dynamics with less recurrent, less diverse, less connected and more segregated phase-synchronization patterns than for conscious states. Using whole-brain models constrained with healthy and injured connectomes, we showed that both pathological and pharmacological low-level states of consciousness presented altered network interactions and more homogeneous and anatomically-constrained local dynamics than conscious states.

It has been proposed that an imbalance between integration and segregation of information in the brain is a network effect of the loss of consciousness [34, 17] that impairs the neural communication across specialized brain modules or subnetworks [35, 36, 37]. Consistent with this view, we found an alteration of integration-segregation of functional phase interactions during low-level states of consciousness caused by brain damage, deep anaesthesia, and anaesthesia’s long-lasting effects during recovery (Fig. 1). Previous work argued that integration and segregation are reconciled in the case of metastability, which has been shown to produce transient synchronized clusters for which sets of brain regions engage and disengage in time, facilitating the exploration of a larger dynamical repertoire of the brain [26, 30, 38]. Here, we showed that the diversity of phase synchronization patterns and their recurrence in time were also reduced in low-level states of consciousness (Fig. 1), presumably leading to a failure to dynamically balance integration and segregation. These results are in line with previous studies showing differences in the synchronized states both in space and time during altered states of consciousness [39, 40, 41, 24, 42, 17, 14, 18].

The whole-brain model used here allowed us to understand how structural, dynamical, local and network properties interplay in the different levels of consciousness. Within this model, the network dynamics depended on three ingredients: local bifurcation parameters, the global strength of connections and their structure. Consistent with previous studies [13, 43], we showed that the brain dynamics of low-level states of consciousness were more constrained by the structural connectivity. In the model, this effect arises due to a reduced global coupling strength in low-level states of consciousness, restricting the propagation of activity to direct connections. In contrast, during conscious wakefulness, sufficient global connectivity allows the propagation of activity through direct and indirect paths, thus enhancing the communication between different brain regions. This result supports the predictions of integrated information theory (IIT), which proposes that unconscious states are characterized by a loss of information propagation and integrative capacity of the brain [35]. The observed decrease in global connectivity is also consistent with previous studies of EEG signals after a transcraneal magnetic stimulation (TMS)-mediated perturbation, showing that low-level states of consciousness were less responsive than conscious states[19, 20].

We studied different versions of the model, which could be homogeneous (all local parameters were constant) or heterogeneous (local parameters were allowed to vary from one brain region to the other and were estimated from the data). Using the heterogeneous model, we found that in low-level states of consciousness, the estimated local dynamics were strongly determined by the structural connections. In contrast, local dynamics associated with consciousness presented a diversity across nodes that were not fully determined by structural connections. In other words, for conscious states, local dynamics can dissociate from their structural constrains, allowing for additional heterogeneity arising from dynamics. Moreover, including the damaged structural connectivities due to brain injuries in the DOC patients into the model showed a further limitation of the diversity of local dynamics in pathological low-level states of consciousness.

These results are consistent with dynamics tied to the structure during low-level states of consciousness and have important functional implications. Indeed, electrophysiological, fMRI and MEG studies have shown that heterogeneous local dynamics, differing between sensory and association brain regions, contribute to the hierarchical specialization across areas at the functional level [44, 45, 46, 47, 48]. Recently, it has been shown that extending models to include heterogeneous information of local dynamics, e.g., as given by positron-emission tomography (PET) maps of neurotransmitter receptor density [46] or by *T_w_1/T_w_2* maps as proxies of microcircuit properties [47], increases model performance to fit empirical data. Our model could be extended to include these and other axes of hierarchy to explore brain mechanism of consciousness.

Furthermore, in parallel with the additional dynamic-based heterogeneity observed in conscious states, we found that local dynamics of hub regions were more stable in conscious states than in low-level states of consciousness and contribute the most to the system’s linear stability (Supplementary Fig. 7). This suggests that in order to release the structural constraints on local dynamics while ensuring the global stability of the system, hubs play an important role by increasing their local stability and diminishing their variability. We believe that the dynamical stability of the hubs is a signature of consciousness and has functional implications. On one hand, unstable hubs would propagate noise to the rest of the network, thus degrading the communication among brain regions. On the other hand, the stability of hubs is required to maintain a functional core-periphery architecture. It has been shown that this architecture is essential to achieve trade-offs between stability and flexibility [49]. Indeed, previous studies of complex systems have derived general principles of core-periphery architecture, pointing that the network periphery can support more variability, responsitivity and plasticity than the network core, while the latter enhances the system robustness [49, 50]. Consistent with this, previous works on whole-brain fMRI have observed core-periphery organization during resting state [51] and a stable core together with a variable periphery during learning [52]. Finally, we showed that structural breakdown of core-periphery architecture, as observed in injured structural connectivity (Fig. 5), also leads to a reduction of dynamical heterogeneity. Thus, functional disruption in low-level states of consciousness might partly rely on an attenuation of core-periphery structure induced by i) the loss of stability of the hubs and ii) the structural damage of the hubs.

Overall, our results suggest that, during healthy wakefulness, in order to allow a dynamically-based heterogeneity of local dynamics across the brain, resulting in diverse collective activity patterns, while preserving stability and a core-periphery architecture, the hubs are required to “anchor” the dynamics by increasing their stability.

A prediction of our study is thus that, under localized external stimulation, hub regions should be less responsive for conscious states compared to low-level states of consciousness. In particular, we found stronger effects in subcortical areas, such as thalamus and hippocampus, and in the precuneus and the posterior cingulate areas, which are directly involved in the thalamo-cortical loop and are thought to down-regulate the activity of the cortical network [53]. Thus, enhancement of neural excitability in those regions through therapeutic procedures may improve conscious recovery process [54]. However, current stimulation protocols using TMS to investigate the network response during different states of consciousness in humans [19, 20] cannot achieve the required localization of stimulation to test our predictions. Indeed, TMS is a strong external perturbation that activates several cortical and subcortical areas, producing a global perturbation. Nevertheless, at the moment, in-silico perturbation of diverse computational models [55, 56] might be useful to test this prediction.

Using global synchronization measures, we found significant differences for different levels of consciousness (CNT and DOC patients and W, S, and R), but these measures mostly failed to identify a significant difference between patients groups (MCS vs. UWS) (Fig. 1). However, our model-based analysis of local dynamics was able to distinguish between patients groups (Figs. 3f, 4, 5h-i and Supplementary Fig. 7). This highlights the clinical translation potential of multi-parameter whole-brain models and the need of further studies that consider region-specific measures for clinical predictions. Nevertheless, given the patient inclusion criteria used here (see Methods), a limitation of our study is the potential lack of generalizability of the results to a broader spectrum of DOC patients, such as those presenting larger brain structural damage.

In conclusion, our results show that pathological and pharmacological low-level states of consciousness presented altered network interactions, more homogeneous, structurally-constrained local dynamics, and less stability of the network’s core compared to conscious states. These results provide relevant information about the mechanisms of consciousness both from the research and clinical point of view.

## 4 Methods

### Participants

In this study, we have selected altered states of consciousness for pathological condition so-called DOC, and healthy subjects during propofol anaesthesia-induced loss of consciousness. The study was approved by the Ethics Committee of the Faculty of Medicine of the University of Liege. Written informed consent to participate in the study was obtained directly from healthy control participants and the legal surrogates of the patients.

We selected 48 DOC patients, 33 in MCS (9 females, age range 24-83 years; mean age ± SD, 45 ± 16 years) and 15 with UWS (6 females, age range 20-74 years; mean age ± SD, 47 ± 16 years) and 35 age and gender-matched healthy controls (14 females, age range 19-72 years; mean age ± SD, 40 ± 14 years). The DOC patients data was recorded 880 ± 35 days after injury. The healthy controls data was collected while awake and aware. The diagnosis of the DOC patients was confirmed through repeated behavioural assessment with the Coma Recovery Scale-Revised (CRS-R) that evaluates auditory, visual, motor, sensorimotor function, communication and arousal [57]. The DOC patients were included in the study, if MRI exam was recorded without anesthetized condition and the behavioural diagnosis was carried out at least five times for each patient using CRS-R examination [58]. The best CRS-R result was retained for the behavioural diagnosis. The exclusion criteria of patients were as follows: (i) having any significant neurological, neurosurgical or psychiatric disorders prior to the brain insult that lead to DOC, (ii) having any contraindication to MRI such as electronic implanted devices, external ventricular drain, and (iii) being not medically stable or large focal brain damage, i.e. > 2/3 of one hemisphere. Details on patients’ demographics and clinical characteristics are summarized in Supplementary Table 7-8.

For the propofol anaesthesia, 16 healthy control subjects (14 females, age range, 18-31 years; mean age ± SD, 22 ± 3.3 years) were selected in three clinical states including normal wakefulness with eyes closed (W), anaesthesia-induced reduction of consciousness (S) and recovery from anaesthesia (R). Propofol was infused through an intravenous catheter placed into a vein of the right hand or forearm and arterial catheter was placed into the left radial artery. During the study ECG, blood pressure, SpO2 and breathing parameters were monitored continuously. The level of consciousness was evaluated clinically throughout the Ramsay scale, representing the verbal commands; for details on the procedure, see [60]. It should be noted that during the recovery of consciousness, R, subjects showed clinical recovery of consciousness (i.e., same score on Ramsay sedation scale as during wakefulness) but they showed residual plasma propofol levels and lower reaction times scores. The healthy subjects did not have MRI contradication, any history of neurological or psychiatric disorders or drug consumption, which have significant effects in brain function.

### MRI acquisition and data analysis

For the DOC dataset, structural and fMRI data were acquired on a Siemens 3T Trio scanner (Siemens Inc, Munich, Germany); propofol dataset was acquired on a 3T Siemens Allegra scanner (Siemens AG, Munich, Germany). The acquisition parameters are described in the Supplementary Information.

The preprocessing of MRI data and the extraction of BOLD time series are described in Supplementary Information. Briefly, independent component analysis was used for motion correction, spatial smoothing and non-brain removal. After preprocessing, FIX (FMRIB’s ICA-based X-noiseifier) [61] was applied to remove the noise components and the lesion-driven artefacts, independently and manually, for each subject. Finally, FSL tools were used to obtain the BOLD time series of 214 cortical and subcortical brain regions (see details in Supplementary material Table 9) in each individual’s native EPI space, defined according to a resting-state Shen atlas [62] and removing the cerebellar parcels.

### Structural connectivity

A whole-brain structural connectivity (SC) matrix was computed for each subject from the DOC dataset, using diffusion imaging and probabilistic tractography (see Supplementary Information for details). The procedure resulted in a symmetric SC matrix summarizing the density of anatomical links among the 214 ROIs, for each healthy control and participant.

### Phase-interaction matrices

To evaluate the level of synchrony in the fMRI the phase interaction between BOLD signals was evaluated. Therefore, a band-pass filter within the narrowband of 0.04 — 0.07 Hz was applied in order to extract the instantaneous phases *φ_j_* (*t*) for each region j. This frequency band has been mapped to the gray matter and captures more relevant information than other frequency bands in terms of brain function [32]. The instantaneous phases, *φ_j_*(*t*), were then estimated applying the Hilbert transform to the filtered BOLD signals individually. The Hilbert transform derives the analytic representation of a real-valued signal given by the BOLD timeseries. The analytical signal, *s*(*t*), represents the narrowband BOLD signal in the time domain. This analytical signal can be also described as a rotating vector with an instantaneous phase, *φ*(*t*), and an instantaneous amplitude, *A(t)*, such that *s*(*t*) = *A*(*t*)*cos*(*φ*(*t*)). The phase and the amplitude are given by the argument and the modulus, respectively, of the complex signal *z*(*t*) = *s*(*t*) + *i.H*[*s*(*t*)], where *i* is the imaginary unit and *H*[*s*(*t*)] is the Hilbert transform of *s*(*t*).

The synchronization between pairs of brain regions was characterised as the difference between their instantaneous phases. At each time point, the phase difference *P_jk_*(*t*) between two regions *j* and *k* was calculated as:

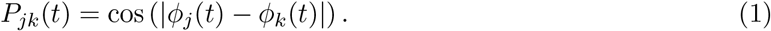

Here, *P_jk_* = 1 when the two regions are in phase (*φ_j_* = *φ_k_*), *P_jk_* = 0 when they are orthogonal and *P_jk_* = —1 when they are in anti-phase. At any time *t*, the phase-interaction matrix *P*(*t*) represents the instantaneous phase synchrony among the different ROIs. The time averaged phase-interaction matrix, 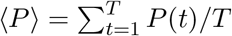, was bias-corrected by subtracting the expected phase-interactions phase-randomized surrogates, designed to decorrelate the phases while preserving the power spectrum of the original signals (see Supplementary Information).

The instantaneous global level of synchrony of the whole network *r*(*t*) was calculated as the average of the phase differences at each time point. Since *P*(*t*) is a symmetric matrix, then:

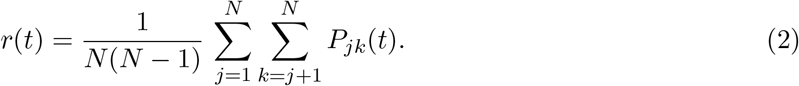

Finally, the fluctuations of *r*(*t*) over time indicate the diversity of the observed network phase inter-actions. The *phase-interaction fluctuations m* were thus calculated as the standard deviation of *r*. When all the nodes of a network are synchronised then *r*(*t*) = 1 for all *t* and thus *m* = 0. However, if the network switches among synchronization states over time leading to fluctuations of *r*, then *m* > 0, reflecting those fluctuations.

### Integration

We used the integration measure to evaluate the brain’s capacity to link network communities and ensure communication. The integration, *I*, was determined using the length of the largest connected component of the time-averaged phase-interaction matrix, 〈*P*〉, based on the procedure presented in [8]. The number of nodes within the largest connected component of the binarized phase-interaction matrix was computed for different binarizing thresholds, ranging from 0 to 1 (scanning the whole range). The largest connected component was given by the largest sub-graph in which any two vertices are connected to each other by paths, and which connects to no additional vertices in the super-graph. We define the integration value, *I*, as the integral of the size of the largest connected component as a function of the threshold.

### Segregation

Complementary to the integration, we measured the brain’s ability to distinguish densely connected network communities. This was done by measuring the segregation of phase-interactions using a community analysis detection. First, we binarized the matrix 〈*P*〉 by detecting the pairs of regions with average phase interaction significantly (*p* < 0.01) larger than expected in phase-randomized surrogates (see Supplementary Information). The segregation was measured in the binarized phase-interaction matrix. It was given by the statistics of the quality of the partition algorithm, i.e., the cost function of the process of detecting communities, or modularity index *Q* [63]. Communities were detected using the Louvain algorithm that performs a subdivision of the matrix into non-overlapping groups of brain regions which maximize the number of within-group edges and minimizes the number of between-group edges [64]. The modularity index, *Q*, measures the statistics of the community detection and evaluates the quality of the partition in terms of the number of within- and between-groups’ edges.

### Functional connectivity dynamics (FCD)

We evaluated the presence of repeating patterns of network states by calculating the recurrence of the phase-interaction patterns. For this, we used the functional connectivity dynamics (FCD). This measure is based on previous studies that defined the FCD for FC matrices calculated in different time windows [65]. In our study, the duration of scans (10 min) was divided into M=20 sliding windows of 30 s, shifted in 2 s steps. For each time window, centred at time *t*, the average phase-interaction matrix, 〈*P*(*t*)〈, was calculated as:

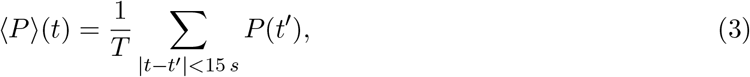

where *T* is the total number of TRs in 30 s. We then constructed the *M* × *M* symmetric matrix whose (*t*_1_, *t*_2_) entry was defined by the cosine similarity, *θ*, between the upper diagonal elements of two matrices 〈*P*〉(*t*_1_) and 〈*P*〉(*t*_2_), given as:

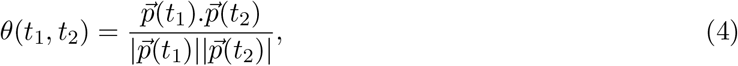

where 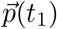 and 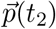 are the vectorized representations of matrices 〈*P*〉(*t*_1_) and 〈*P*〉(*t*_2_), respectively. Finally, the FCD measures was given by the distribution of these cosine similarities for all pairs of time windows.

### Whole-Brain Network Model

The brain network model consists of *N* = 214 coupled brain regions derived from the Shen parcellation [62]. The global dynamics of the brain network model used here results from the mutual interactions of local node dynamics coupled through the underlying empirical anatomical structural connectivity matrix *C_ij_* (see Fig. 1). Local dynamics are simulated by the normal form of a supercritical Hopf bifurcation, i.e., Stuart-Landau oscillator [66, 67], describing the transition from noisy oscillations to sustained oscillations [68], and is given, in the complex plane, as:

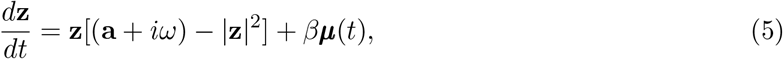

where *Z_j_* is a complex number, *μ_j_*(*t*) is additive Gaussian noise with standard deviation *β* = 0.02, and *ω_j_* corresponds to the intrinsic frequency of the oscillator in the range of 0.04-0.07 Hz band. The intrinsic frequencies were estimated from the averaged peak frequency of the narrowband empirical BOLD signals of each brain region. For *a_j_* < 0, the local dynamics present a stable spiral point, producing damped or noisy oscillations in absence or presence of noise, respectively (Supplementary Fig. 2). For *a_j_* > 0, the spiral becomes unstable and a stable limit cycle oscillation appears, producing autonomous oscillations with frequency 2*πf_j_* = *W_j_* (Supplementary Fig. 2). The BOLD fluctuations were modelled by the real part of the state variables, i.e., Real(*z_j_*).

The whole-brain dynamics were obtained by coupling the local dynamics through the *C_ij_* matrix:

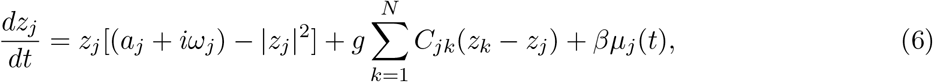

where *g* represents a global coupling scaling the structural connectivity *C_i_j*. The matrix *C_i_j* is scaled to a maximum value of 0.2 to prevent full synchronization of the model. Interactions were modelled using the common difference coupling, which approximates the simplest (linear) part of a general coupling function [69].

### Homogeneous model: Fitting the global coupling *g*

To create a representative model of BOLD activity in each brain state, we adjusted the model parameters (*g* and *a_j_*) to fit the spatiotemporal BOLD dynamics for each brain state and each dataset.

Our first aim was to describe the global properties of the spatio-temporal dynamics of each subject in each state, independently of the variations in the dynamics of local nodes. For that reason, in this first approach to the model, all nodes were set to *a_j_* = 0, called the homogeneous model. The global coupling parameter *g* was obtained by fitting the simulated and empirical data. Specifically, for each value of *g*, the model FCD was computed and compared with the empirical FCD using the Kolmogorov-Smirnov (KS) distance between the simulated and empirical distribution of the FCD elements. The KS-distance quantifies the maximal difference between the cumulative distribution functions of the two samples. Thus, the optimal value of *g* was the one that minimized the KS distance.

### Heterogeneous model: Local optimization of the bifurcation parameters

To evaluate the heterogeneous local dynamics on the network’s dynamics, we extended the model to allow differences in bifurcation parameters *a_j_* for different ROIs. The *g* parameter was the one estimated with the homogeneous model. The bifurcation parameters were optimized based on the empirical power spectral density of the BOLD signals in each node. Specifically, we fitted the proportion of power in the 0.04-0.07 Hz band with respect to the 0.04-0.25 Hz band (i.e. we removed the smallest frequencies below 0.04 Hz and considered the whole spectrum up to the Nyquist frequency which is 0.25Hz) [33]. For this, the BOLD signals were filtered in the 0.04-0.25 Hz band and the power spectrum *PS_j_*(*f*) was calculated for each node *j*. We then defined the proportion of power in the 0.04-0.07 Hz band as:

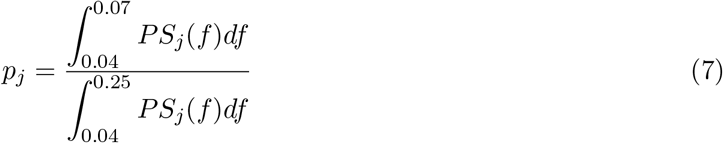

We updated the local bifurcation parameters by an iterative gradient descendent strategy, i.e.:

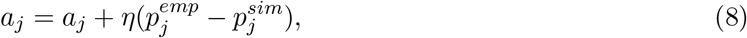

until convergence. *η* was set to 0.1 and the updates of the *a_j_* values were done in each optimization step in parallel.

### Relation between the weight of the strength of a node and its dynamics

Finally, the relation between local and network dynamics was studied. An effective bifurcation parameter 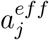 was defined which contains information of the local dynamics and local structure given by its strength. This parameter permits to extract the relation between the dynamics and structure of each node. More specifically, in equation 6, we separated the part that relates to the effective local dynamics and the part that relates to the interaction between nodes. Noting that 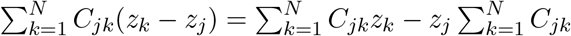, equation 6 can be written as:

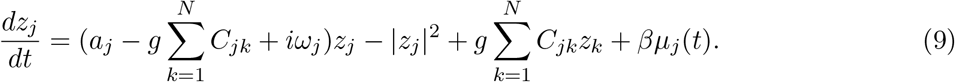

Taking 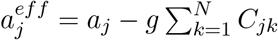, we get:

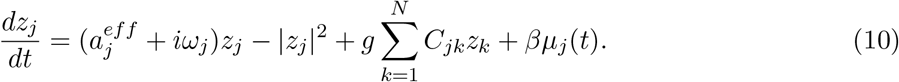

Note that, if *a_j_* is homogeneous across the network (*a_j_* = *a* for all *j*), 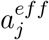 is linearly related to the nodal strength 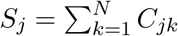.

### Graph analysis of the structural connectivity

The network organization of the SC matrices was investigated using measures of graph theory (GAlib: Graph Analysis library in Python / Numpy, www.github.com/gorkazl/pyGAlib). Here, we focused only on the identification of hub regions and a rich-club to study their relation to the dynamical properties. The strength of a node is the number of connections a node makes in a network. Hubs are thus defined as nodes with high strength, usually playing a central role in the network’s communication. Computing the rich club coefficients requires that the weighted SCs derived from tractography are binarized, discarding the smaller values. An adaptive threshold was applied such that all resulting binary SCs had the same number of links, with a link density of 0.2. A rich club is a supra-structure of a network happening when (i) a network contains hubs and (ii) those hubs are densely connected with each other, forming a cluster [70]. Identifying the presence or the absence of a rich-club is a sensitive problem because it relies on the interpretation of an indirect metric, *k*-density, *ρ*(*k*), an iterative process which evaluates the density *ρ*(*k*’) of the network after all nodes with degree *k* < *k*’ have been removed [70]. Here, we considered a strict criterion and considered that networks contained a rich-club only if a degree *k*’ exist for which *ρ*(*k*’) overcomes 0.9 (largest possible density is 1.0), implying that the hubs of the network are almost all-to-all connected. The regions forming the rich-club were thus identified as the remaining set of nodes with degree *k* > *k*’. Finally, to study alterations in the rich-club due to brain damage, rich-club identification was performed from the SCs of all patients, all healthy controls and from the averaged SC for the control group. The probability of a brain region to take part in the rich-club was evaluated as the frequency with which the region is present in the rich-clubs identified across subjects of the same group.

### Statistical analysis

Statistical differences between levels of consciousness were assessed using one-way repeated measures (rm) ANOVA followed by multiple comparisons using False Discovery Rate (FDR) correction [71]. The threshold for statistical significance was set to p-values<0.05. Wilcoxon rank-sum test (equivalent to a Mann-Whitney U test) was applied in order to find region-wise differences between CNT and DOC patients in the strength of the SC. We corrected for multiple comparisons by using the FDR correction, considering P<0.05 as statistically significant.

## Data availability

The codes and multimodal neuroimaging data from the experiment are available upon request.

## Authors’ contributions

ALG, APA, GZL, SL and GD designed research. RP, JA, AT, CM and OG acquired the data. ALG, RP, AE and MK preprocessed the data. ALG and RJ analysed the data. ALG, APA, GZL, MK and GD studied the computational model. APA, GZL, OG, SL and GD supervised research. ALG, RP, APA, AE and GZL wrote the manuscript. All authors contributed to the editing of the manuscript.

## Acknowledgements

ALG and GD was supported by Swiss National Science Foundation Sinergia grant no. 170873. APA and GD received funding from the FLAG-ERA JTC (PCI2018-092891). GD acknowledges funding from the European Union’s Horizon 2020 FET Flagship Human Brain Project under Grant Agreement 785907 HBP SGA1, SGA2 and SGA3, the Spanish Ministry Project PSI2016-75688-P (AEI/FEDER), the Catalan Research Group Support 2017 SGR 1545, and AWAKENING (PID2019-105772GB-I00, AEI FEDER EU) funded by the Spanish Ministry of Science, Innovation and Universities (MCIU), State Research Agency (AEI) and European Regional Development Funds (FEDER). The study was further supported by the University and University Hospital of Liege, the Belgian National Funds for Scientific Research (FRS-FNRS), the European Space Agency (ESA) and the Belgian Federal Science Policy Office (BELSPO) in the framework of the PRODEX Programme, “Fondazione Europea di Ricerca Biomedica”, the Bial Foundation, the Mind Science Foundation and the European Commission, the fund Generet, the King Baudouin Foundation, AstraZeneca foundation, and the DOCMA project [EU-H2020-MSCA-RISE-778234]. RP is research fellow. OG is research associate and SL is research director at F.R.S.-FNRS. MLK is supported by the ERC Consolidator Grant: CAREGIVING (n. 615539), Center for Music in the Brain, funded by the Danish National Research Foundation (DNRF117), and Centre for Eudaimonia and Human Flourishing funded by the Pettit and Carlsberg Foundations.

We would like to thank the healthy participants and the patients, their families, caregivers and treating clinicians for their participation in this study. The authors thank the whole staff from the ICU and Nuclear Medicine departments, University Hospital of Liège. We are highly grateful to the members of the Liege Coma Science Group for their assistance in clinical evaluations. We also thank Manel Vila-Vidal for his valuable comments.

