## Supplementary Information for "Loss of consciousness reduces the stability of brain hubs and the heterogeneity of brain dynamics"

### 1 Methods

#### 1.1 MRI acquisition and data analysis

For the DOC dataset, structural and functional MRI (fMRI) data were acquired on a Siemens 3T Trio scanner (Siemens Inc, Munich, Germany). The BOLD fMRI resting state (i.e. task free) was acquired using EPI, gradient echo with following parameters: volumes = 300, TR = 2000 ms, TE = 30 ms, flip angle = 78°, voxel size = $3 \times 3 \times 3$ mm$^3$, FOV = $192 \times 192$ mm$^2$, 32 transversal slices, with a duration of 10 minutes. Subsequently, structural 3D T1-weighted MP-RAGE images with were acquired with following parameters: 120 transversal slices, TR = 2300 ms, voxel size = $1.0 \times 1.0 \times 1.2$ $mm^3$, flip angle = 9°, FOV = $256 \times 256$ mm$^2$. Last, diffusion weighted MRI (DWI) was acquired in 64 directions (b-value =1,000 s/mm$^2$, voxel size = $1.8 \times 1.8 \times 3.3$ mm$^3$, FOV = $230 \times 230$ mm$^2$, TR= 5,700 ms, TE= 87 ms, 45 transverse slices, $128 \times 128$ voxel matrix) preceded by a single unweighted image (b0). The DWI was acquired twice.

The propofol dataset was acquired on a 3T Siemens Allegra scanner (Siemens AG, Munich, Germany). The fMRI resting state were acquired using the following parameters: EPI, gradient echo, volumes = 200; TR = 2460ms, TE = 40 ms, voxel size = $3.45 \times 3.45 \times 3$ mm$^3$, FOV = $220 \times 220$ mm, 32 transverse slices, $64 \times 64 \times 32$ matrix size. The structural images were acquired using 3D T1-weighted MP-RAGE with following parameters: 120 transversal slices, TR = 2250 ms, TE=2.99ms, voxel size = 1 mm3, flip angle = 9°, FOV = $256 \times 240 \times 160$mm.

Preprocessing of MRI data was performed using MELODIC (Multivariate Exploratory Linear Optimized Decomposition into Independent Components) version 3.14 [1], which is part of the FMRIB's Software Library (FSL, `http://fsl.fmrib.ox.ac.uk/fsl`). Preprocessing steps included: discarding the first 5 volumes, motion correction using MCFLIRT [2], non-brain removal using BET (Brain Extraction Tool) [3], spatial smoothing with 5 mm FWHM Gaussian Kernel, rigid-body registration, high pass filter cutoff = 100.0 s, and single-session ICA with automatic dimensionality estimation. After preprocessing, FIX (FMRIB's ICA-based X-noiseifier) [61] was applied to remove the noise components and the lesion-driven artefacts, independently, for each subject. Specifically, FSLeyes package in Melodic mode was used to manually classify the single-subject Independent Components (ICs) into "good" for signal, "bad" for noise or lesion-driven artefacts and "unknown" for ambiguous components. Each component was classified by looking at the spatial map, the time series, and the temporal power spectrum [4, 5]. Finally, FIX was applied by using the default parameters to obtain a cleaned version of the functional data.

FSL tools were used to obtain the blood-oxygen-level-dependent (BOLD) time series of the 214 cortical and subcortical brain regions (without the cerebellum, see more details in Supplementary material Table S7) in each individual's native EPI space, defined according to a resting-state atlas [62]. Specifically, the cleaned functional data previously obtained were co-registered to the T1-weighted structural image by using FLIRT [6]. Then, the T1-weighted image was co-registered to the standard MNI space by using FLIRT (12 DOF) and FNIRT [6, 7]. The resulting transformations were concatenated and inverted and applied to warp the resting-state atlas from MNI space to the cleaned functional data. To ensure the preservation of the labels, a nearest-neighbour interpolation method was used. Then, the BOLD time series for each of the 214 brain regions were extracted for each subject in their native space by using *fslmaths* to obtain a binary mask of each brain region, and *fslmeants* to obtain the time series of each binary mask.

The grand average of the functional connectivity matrix, FC, was constructed using Matlab 2017 (The MathWorks Inc.) to compute the pairwise Pearson correlation between all 214 brain regions, applying Fisher's transform to the r-values to get the z-values for the final $214 \times 214$ functional connectivity matrices.

#### 1.2 Structural connectivity

A whole-brain structural connectivity (SC) matrix was computed for each subject from the DOC dataset, using two-step process as described in previous studies [8, 9, 10]. Similar to the procedure used for analysing the resting-state fMRI data, we used the resting-state atlas to create a structural connectome in each individual's diffusion native space. First, DICOM images were converted to Neuroimaging Informatics Technology Initiative (NIfTI) format using dcm2nii (`www.nitrc.org/projects/dcm2nii`). The b0 image in DTI native space was co-registered to the T1-weighted structural image by using FLIRT [6]. The T1-weighted structural image was co-register to the standard space by using FLIRT and FNIRT [6, 7]. The resulting transformations were inverted and applied to warp the resting-state atlas from MNI space to the native MRI diffusion space by applying a nearest-neighbour interpolation algorithm. Second, analysis of diffusion images was performed using the processing pipeline of the FMRIB's Diffusion Toolbox (FDT) in FMRIB's Software Library (`www.fmrib.ox.ac.uk/fsl`). The non-brain tissues were extracted by applying the Brain Extraction Tool (BET) [3], the eddy current distortions and head motion were corrected using eddy correct tool [11], and the gradient matrix was reoriented to correct for subject motion [12]. Then, Crossing Fibres were modelled using the default BEDPOSTX parameters and the probability of multi-fibre orientations were calculated to improve the sensitivity of non-dominant fibre populations [13, 14]. Then, Probabilistic Tractography was performed in native MRI diffusion space using the default settings of PROBTRACKX [13, 14]. For each brain region, the connectivity probability to each of the other 213 brain regions was computed. The resulting matrix was then symmetrized by computing their transpose matrix and averaging both matrices, therefore $C_{ij} = C_{ji}$. Finally, to obtain the structural probability matrix, the value of the probability pairs of brain regions was divided by its corresponding number of generated tracts. To summarize, for each participant, a 214x214 symmetric weighted network was constructed and normalised by the total number of fibres in the whole network; thus, the structural connectivity matrix (SC) represents the density of links of the anatomical organization of the brain.

#### 1.3 Surrogate Analysis

For the phase surrogate analysis, first, the Fourier transform (FT) of the signals was computed. The phase of the Fourier transform was substituted with uniformly distributed random numbers while preserving their modulus. Then, the inverse FT was applied to return to the time domain with the new Fourier coefficients. This procedure effectively randomizes the phases of the signals while preserving the same power spectra as the original time-courses. Specifically, let $x_i(t)$ be the original BOLD time-course from the brain area $i$. The discrete Fourier transform of $x_i$ is given by:

$$x_i(k) = \sum_{t=1}^{T} x_i(t) e^{-j\frac{2\pi kt}{T}} \tag{1}$$

where $j$ is the imaginary unit and k goes from 1 to T (k=1,...,T). The phase shuffled surrogate is given by:

$$x_i^{surr}(k) = \frac{1}{T}\sum_{t=1}^{T} |x_i(t)| e^{-j(\frac{2\pi kt}{T}+\varphi_r)} \tag{2}$$

where $\varphi_r$ is random variable uniformly distributed between $-\pi$ and $\pi$. These surrogates were used to rerun the analysis and extract a phase interaction matrix used to clean the empirical matrices.

### 1.4 Linear stability analysis

In this section we studied the linear stability of the whole-brain network. The model consists of 214 coupled brain regions, with local Hopf dynamics, coupled through the connectome matrix $\mathbf{C}$. The dynamical system can be written in vector form as:

$$\frac{d\mathbf{z}}{dt} = (\mathbf{a} - \mathbf{S} + i\boldsymbol{\omega}) \odot \mathbf{z} - \mathbf{z} \odot \bar{\mathbf{z}} + G\mathbf{C}\mathbf{z} + \beta\boldsymbol{\mu}(t), \tag{3}$$

where $\odot$ is the Hadamard element-wise product, $\boldsymbol{z} = [z_1, ..., z_N]$ are the complex-valued state variables of each node, $\bar{\boldsymbol{z}}$ is the complex conjugate of $\boldsymbol{z}$, $\mathbf{a} = [a_1, ..., a_N]$ and $\boldsymbol{\omega} = [\omega_1, ..., \omega_N]$ are the vectors containing the bifurcation parameters and intrinsic frequencies of each node, respectively, $\boldsymbol{S} = [S_1, ..., S_N]$ is the vector containing the strength of each node, where $S_j = G\sum_{k=1}^{N} C_{jk}$, and $\boldsymbol{\mu} = [\mu_1, ..., \mu_N]$ is a Gaussian noise vector. The model parameters $\mathbf{a}$, $\boldsymbol{\omega}$, and $G$ were estimated from the data, using the heterogeneous model, for each experimental condition, as described in the main text.

We studied the linear stability of the fixed point $\boldsymbol{z} = \mathbf{0}$, which is solution of $\frac{d\mathbf{z}}{dt} = 0$. In the linearized system the quadratic terms (i.e., $\boldsymbol{z} \odot \bar{\boldsymbol{z}}$) are not taken into account and the evolution of fluctuations $\delta\boldsymbol{z}$ around $\boldsymbol{z} = \mathbf{0}$ can be approximated as:

$$\frac{d}{dt}\delta\mathbf{z} = \mathbf{A}\delta\mathbf{z} + \beta\boldsymbol{\mu}(t), \tag{4}$$

where $\mathbf{A}$ is the Jacobi matrix, given as: $\mathbf{A} = \mathrm{diag}(\mathbf{a} - \mathbf{S} + i\boldsymbol{\omega}) + G\mathbf{C}$, and $\mathrm{diag}(\boldsymbol{x})$ is the diagonal matrix whose entries are the elements of the vector $\boldsymbol{x}$.

The stability of the system is determined by the eigen-decomposition of the Jacobi matrix. Figure S6 shows the eigenvectors of $\mathbf{A}$ as a function of the node strengths and the real part of the eigenvalues associated to the eigenvectors, noted $\mathrm{Real}(\lambda)$. The system is stable since all eigenvalues have $\mathrm{Real}(\lambda) < 0$ and, for all models, the nodes with higher strength, i.e., the network hubs, contribute the most to the most stable eigenvectors, i.e., those with lowest $\mathrm{Real}(\lambda)$. However, the stability of these eigenvectors is reduced for models estimated from data in low-level states of consciousness.

### 2 Supplementary figures

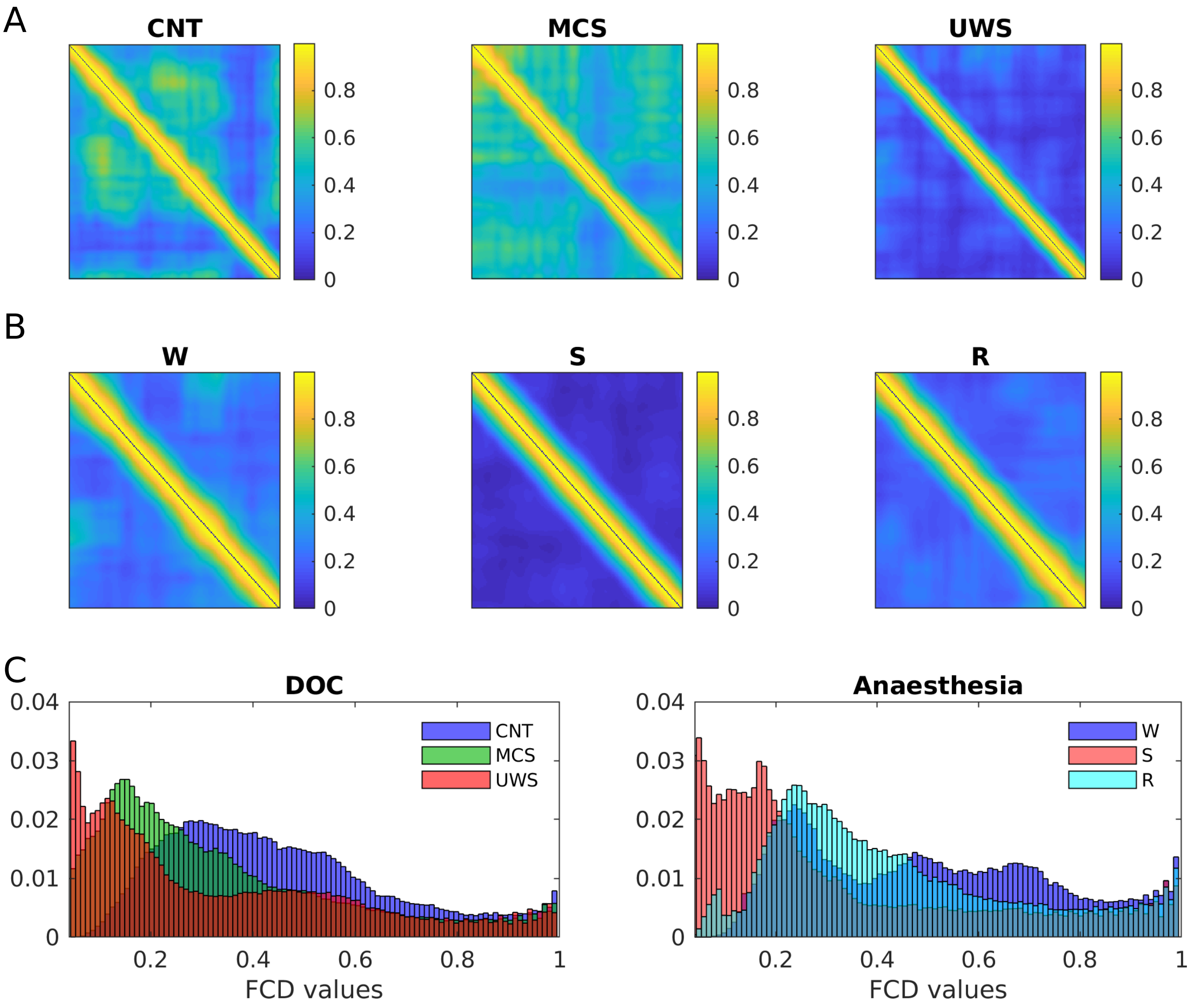


Figure S1: **Functional Dynamical Connectivity (FCD) matrices and distributions of their values.** The FCD matrices were calculated based on the correlation between phase difference matrices at different time windows. **A)** Examples of the FCD for individual healthy subjects and DOC patients. **B)** FCD matrices for individual subjects during wakefulness, deep sedation and recovery from anaesthesia. **C)** The distribution of the upper diagonal elements of the FCD matrix, for each subject group. The distribution of the conscious states (controls and W) was sparser and shifted towards higher values compared to low-level states of consciousness.

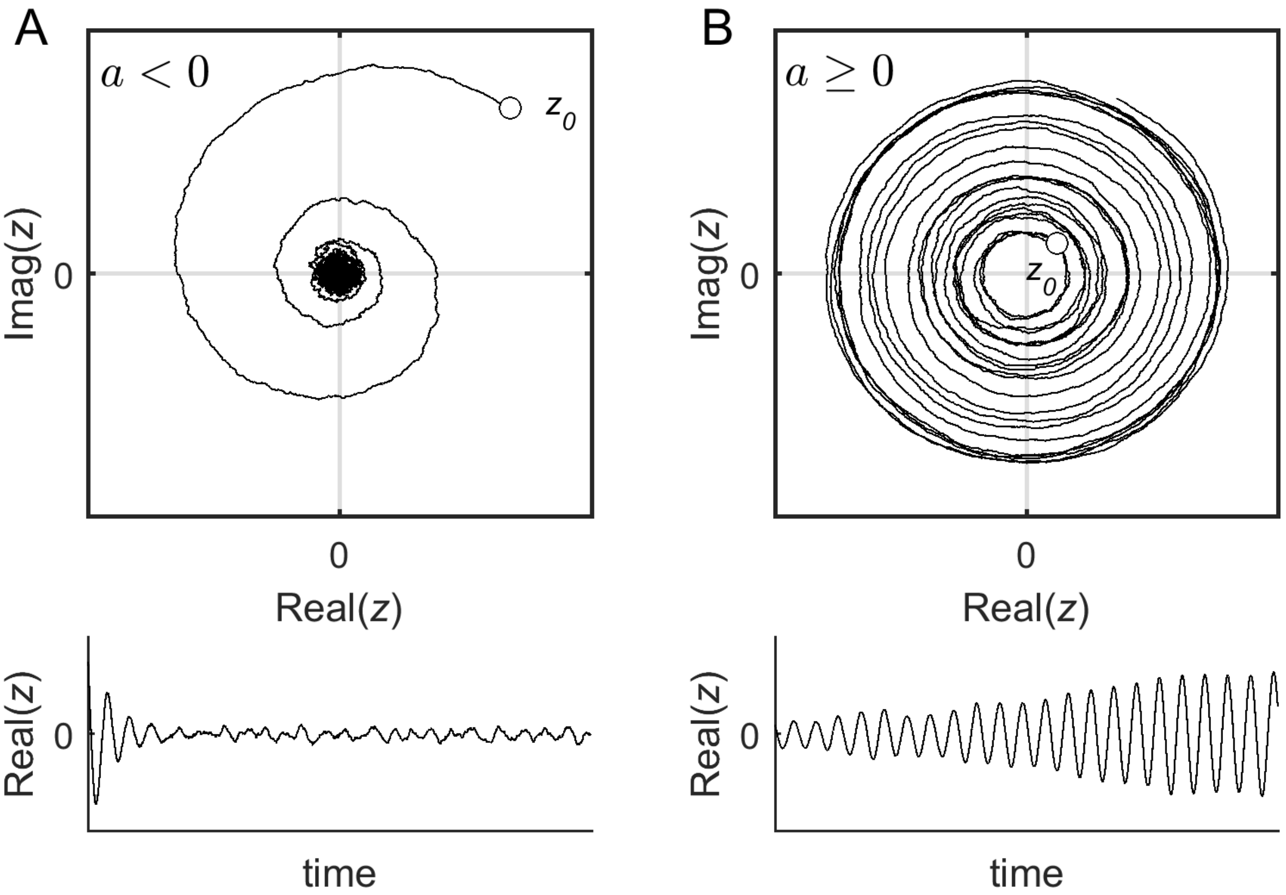


Figure S2: **Phase space for an example of a single Hopf oscillator.** **A)** Subcritical Hopf oscillator ($a < 0$). Top: In this regime, a stable spiral, or focus, exists at $\boldsymbol{z} = 0$. The system relaxes towards the focus with damped oscillations. In the presence of noise, however, the system fluctuates around the focus, thus producing noise-induced oscillations. $\boldsymbol{z_0} = \boldsymbol{z}(t = 0)$ indicates the initial condition. Bottom: temporal evolution of $Real(z)$. **B)** Supercritical Hopf bifurcation ($a \geq 0$). Top: In this regime, the focus at $\boldsymbol{z} = 0$ becomes unstable and a stable limit-cycle appears, thus producing autonomous or self-sustained oscillations. Bottom: temporal evolution of $Real(z)$.

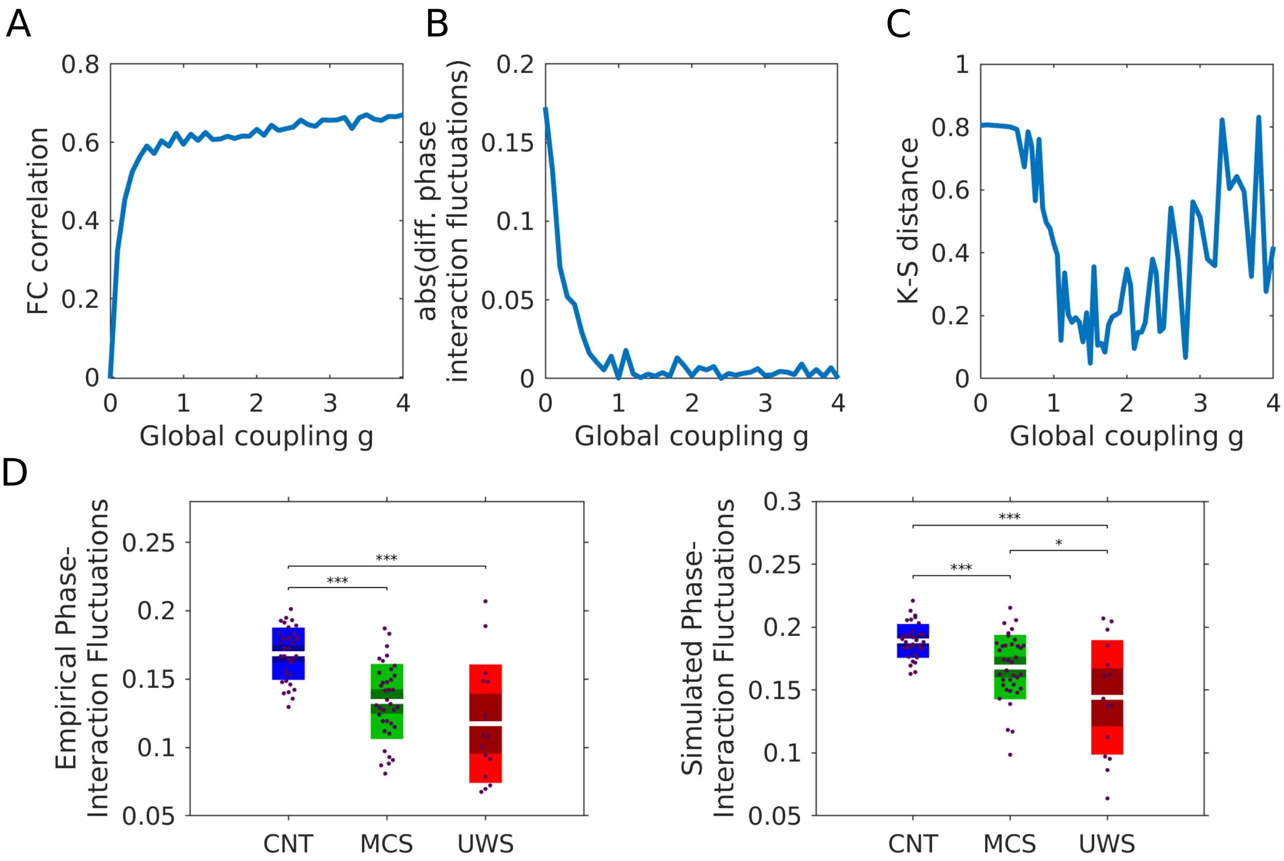


Figure S3: **Goodness of fit of the whole-brain computational model computed by using different measures. A)** Pearson correlation between the empirical and simulated functional connectivity (FC) matrices as a function of parameter $G$. **B)** Absolute difference between the empirical and simulated phase-interaction fluctuations as a function of $G$. **C)** Kolmogorov-Smirnov distance between the empirical and simulated FCD distributions. **D)** The optimal value of $G$ was obtained using the KS-distance between the empirical and simulated FCD distributions. We verified that, for the obtained values of $G$ in each condition, the differences in metastability observed in the data were preserved in the model. For this we used the SC matrix of individual subjects and the parameter $G$ was fixed for each group.

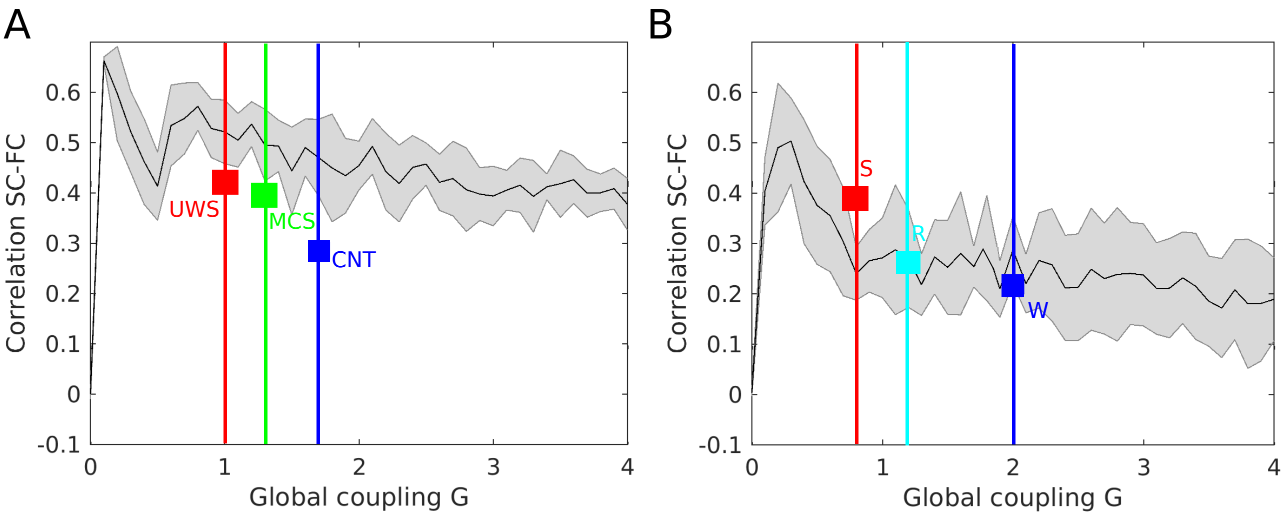


Figure S4: **Empirical and model correlations between the SC and FC matrices for each group.** In **A)** and **B)** the black line corresponds to the correlation between the empirical SC and the FC simulated with the whole-brain model learned from the different experimental groups, as a function of $G$. The shaded area corresponds to the standard error of the values obtained for different simulations. The curve shows a peak for small $G$ and then it decreases slowly as $G$ increases. In **A)** the lines are located in the corresponding optimal global coupling G for each brain state. In **B)** the lines correspond to the states for the anaesthesia dataset. For all cases, the values correspond to the optimal G extracted from the homogeneous model (depicted in Fig. 4). In **A)** and **B)** the squares correspond to the empirical values of the empirical SC and FC correlation. Both empirically and using the model, we observed a shift to smaller values in the SC-FC correlation while the level of consciousness decreases.

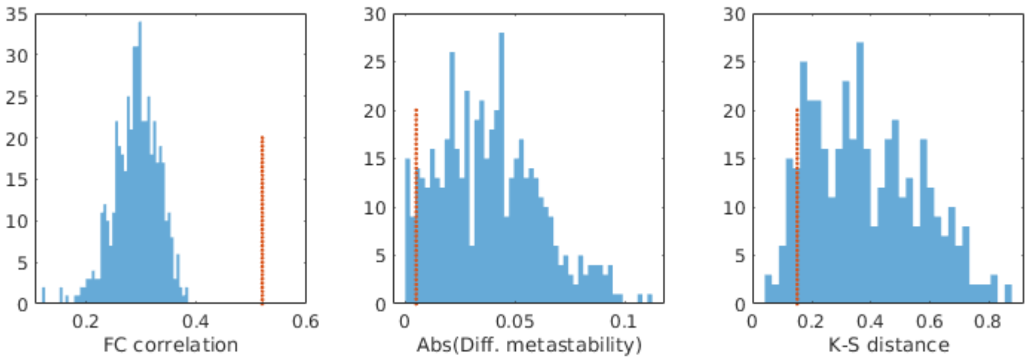


Figure S5: **Goodness of fit of the heterogeneous model compared to models shuffling the order of the a's.** Goodness of fit for the Pearson correlation of the FC matrices, the absolute difference in metastability and the Kolmogorv-Smirnov distance between the FCD matrices. The blue histograms correspond to the fitting after shuffling the labels of the $a_i$ values and the red line corresponds to the mean fitting of the heterogeneous model.

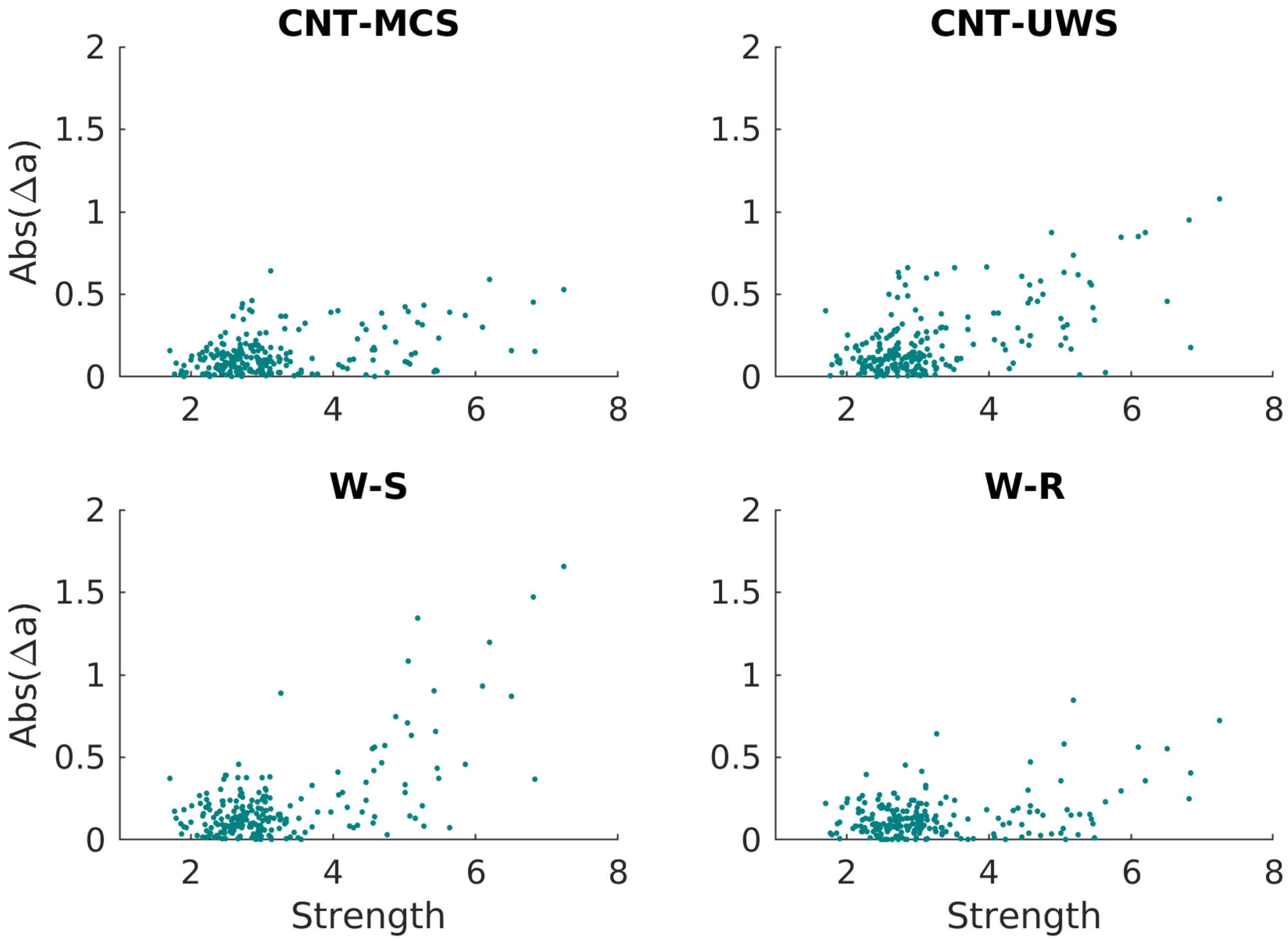


Figure S6: **Relationship between the absolute difference Δa and the strength of each node. A-D)** The absolute difference of the $a$ parameter values between different groups in function of the strength of the nodes extracted from the SC of the healthy controls. Pearson correlations: CNT-MCS: $\rho = 0.40$, ($p < 0.001$); CNT-UWS: $\rho = 0.60$, ($p < 0.001$); W-S: $\rho = 0.63$, ($p < 0.001$); W-R: $\rho = 0.37$, ($p < 0.001$).

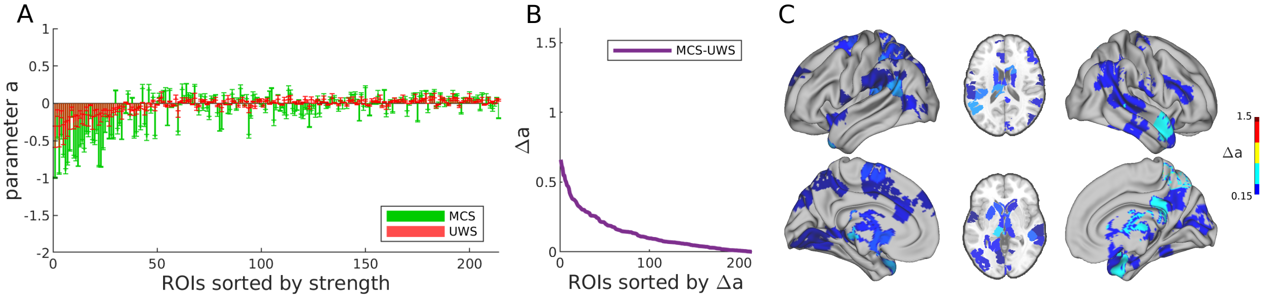


Figure S7: **Comparison between the local bifurcation parameters of the whole-brain model between DOC patients (MCS and UWS). A)** Bars indicate the mean ± standard deviation across simulations of estimated bifurcation model parameters for each of the 214 nodes (sorted by node strength). Green bars correspond to MCS and red bars to UWS. **B** Ranked absolute parameter difference, $\Delta a$, for each the comparison between the two groups. **C)** Spatial distribution of the most altered bifurcation parameter values.

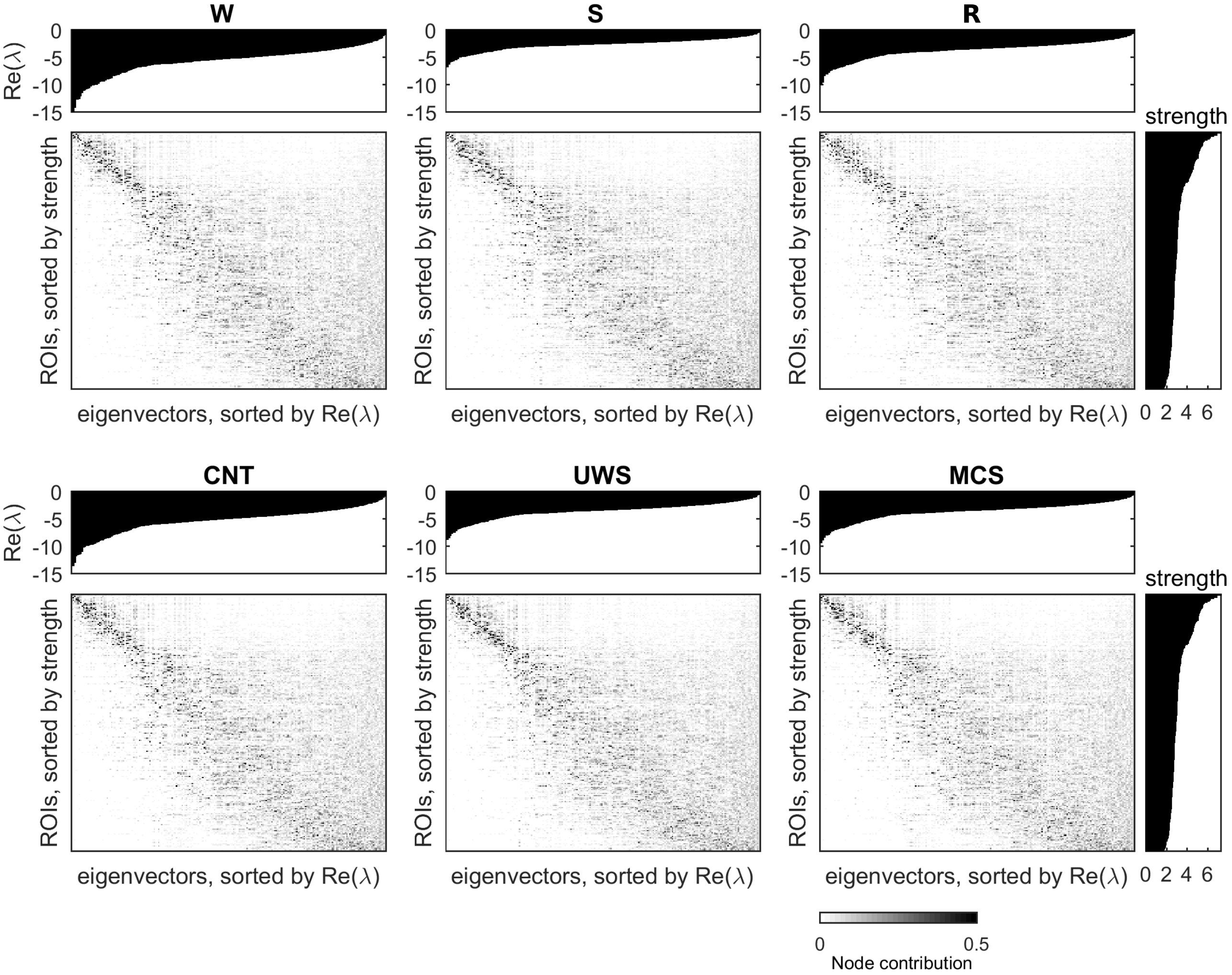


Figure S8: **Eigendecomposition of the Jacobi matrix.** The eigenvectors of the Jacobi matrix ($N$-dimensional vectors) were sorted according to the real part of the associated eigenvalues (top insets), Real($\lambda$), and the strength of the nodes (right insets).

| |
|---|
| 38% Superior frontal gyrus, orbital part R / 25% Middle frontal gyrus, orbital part R |
| 42% Gyrus rectus R / 19% Olfactory cortex R |
| 69% Superior frontal gyrus, orbital part R |
| 44% Middle frontal gyrus R/ 31% Anterior cingulate and paracingulate gyri |
| 34% Insula R/ 22% Lenticular nucleus, putamen R |
| 56% Temporal pole: middle temporal gyrus R/ 20% Fusiform gyrus R |
| 67% Inferior temporal gyrus R |
| 55% Fusiform gyrus R/ 41% Inferior temporal gyrus R |
| 33% Fusiform gyrus R/ 27% Inferior temporal gyrus R |
| 70% Inferior temporal gyrus R |
| 42% Median cingulate and paracingulate gyri R |
| 50% Hippocampus R/ 9% ParaHippocampal gyrus R |
| 30% Hippocampus R/ 15% Amygdala R |
| 41% Thalamus R/ 8% Hippocampus R |
| 52% Thalamus R |
| 40% Superior frontal gyrus, orbital part L/ 34%Middle frontal gyrus, orbital part L |
| 58% Paracentral Lobule L / 35% Precuneus L |
| 61% Inferior temporal gyrus L |
| 55% Temporal pole: superior temporal gyrus L/ 42% Temporal pole: middle temporal gyrus L |
| 45% emporal pole: middle temporal gyrus L/ 25% Fusiform gyrus L |
| 87% Inferior temporal gyrus L |
| 65% Inferior temporal gyrus L |
| 59% Calcarine fissure and surrounding cortex L/ 7% Lingual gyrus L |
| 55% Posterior cingulate gyrus L / 21% Precuneus L |
| 31% Amygdala L / 18%Temporal pole: superior temporal gyrus L |
| 42% Hippocampus L/ 5% Thalamus L |
| 30% Hippocampus L / 21% ParaHippocampal gyrus L |
| 84% Lenticular nucleus, putamen L |
| 53% Thalamus L |
| 51% Thalamus L / 2% Caudate nucleus L |

Table S1: **ROIs with the highest absolute difference in the bifurcation parameters when comparing normal wakefulness in healthy controls and MCS.** All the ROIs showed an absolute difference greater than the threshold given by the sum of the mean and standard deviation of the absolute differences corresponding to all the ROIs. The percentage corresponds to the covered part of the ROI in the Automated Anatomical Labeling (AAL) parcellation code.

| |
|---|
| 42 % Gyrus rectus R / 19 % Olfactory cortex R |
| 69 % Superior frontal gyrus, orbital part R |
| 34 % Insula R/ 22 % Lenticular nucleus, putamen R |
| 56 % Temporal pole: middle temporal gyrus R/ 20 % Fusiform R |
| 67 % Inferior temporal gyrus R |
| 48 % Fusiform R/ 38 % Inferior temporal gyrus R |
| 55 % Fusiform R/ 41 % Inferior temporal gyrus R |
| 33 % Fusiform R/ 27 % Inferior temporal gyrus R |
| 44 % Calcarine fissure and surrounding cortex R/ 6 % Lingual gyrus R |
| 28 % Amygdala R/ 27 % Temporal pole: superior temporal gyrus R |
| 50 % Hippocampus R/ 9 % Parahippocampal gyrus R |
| 30 % Hippocampus R / 15 % Amygdala R |
| 62 % Caudate nucleus R |
| 46 % Caudate nucleus R / 6 % Thalamus R |
| 84 % Lenticular nucleus, putamen R |
| 36 % Thalamus R/ 9 % Lingual gyrus R |
| 41 % Thalamus R / 8 % Hippocampus R |
| 51 % Inferior frontal gyrus, orbital part L / 31 % Superior frontal gyrus, orbital part L |
| 56 % Gyrus rectus L/ 13 % Olfactory cortex L |
| 34 % Insula L/ 18 % Superior temporal gyrus L |
| 61 % Inferior temporal gyrus L |
| 45 % Temporal pole: middle temporal gyrus L/ 25 % Fusiform L |
| 46 % Inferior temporal gyrus L/ 41 % Fusiform L |
| 87 % Inferior temporal gyrus L |
| 65 % Inferior temporal gyrus L |
| 59 % Fusiform L/ 21 % Inferior temporal gyrus L |
| 59 % Calcarine L/ 7 % Lingual gyrus L |
| 55 % Posterior cingulate gyrus L/ 21 % Precuneus L |
| 42 % Hippocampus L/ 5 % Thalamus L |
| 30 % Hippocampus L/ 21 % Parahippocampal gyrus L |
| 28 % Caudate nucleus L/ 15 % Olfactory cortex L |
| 52 % Caudate nucleus L/ 2 % Thalamus L |
| 51 % Thalamus L/ 2 % Caudate nucleus L |

Table S2: **ROIs with the highest absolute difference in the bifurcation parameters when comparing normal wakefulness in healthy controls and UWS.** All the ROIs showed an absolute difference greater than the threshold given by the sum of the mean and standard deviation of the absolute differences corresponding to all the ROIs. The percentage corresponds to the covered part of the ROI in the Automated Anatomical Labeling (AAL) parcellation code.

| |
|---|
| 55 % Fusiform R / 41 % Inferior temporal gyrus R |
| 44 % Calcarine fissure and surrounding cortex R/ 6 % Lingual gyrus R |
| 50 % Hippocampus R / 9 % Parahippocampal gyrus R |
| 30 % Hippocampus R / 15 % Amygdala R |
| 62 % Caudate nucleus R |
| 46 % Caudate nucleus R/ 6 % Thalamus R |
| 57 % Caudate nucleus R/ 11 % Olfactory cortex R |
| 84 % Lenticular nucleus, putamen R |
| 41 % Thalamus R/ 8 % Hippocampus R |
| 83 % Inferior temporal gyrus L |
| 59 % Calcarine fissure and surrounding cortex L/ 7 % Lingual gyrus L |
| 37 % Median cingulate and paracingulate gyri L/ 36 % Anterior cingulate and paracingulate gyri L |
| 55 % Posterior cingulate gyrus L / 21 % Precuneus L |
| 42 % Hippocampus L / 5 % Thalamus L |
| 30 % Hippocampus L/ 21 % Parahippocampal gyrus L |
| 46 % Hippocampus L / 19 % Inferior temporal gyrus L |
| 28 % Caudate nucleus L / 15 % Olfactory cortex L |
| 52 % Caudate nucleus L / 2 % Thalamus L |
| 51 % Thalamus L / 2 % Caudate nucleus L |

Table S3: **ROIs with the highest absolute difference in the bifurcation parameters when comparing normal wakefulness, W and deep sedation, S.** All the ROIs showed an absolute difference greater than the threshold given by the sum of the mean and standard deviation of the absolute differences corresponding to all the ROIs. The percentage corresponds to the covered part of the ROI in the Automated Anatomical Labeling (AAL) parcellation code.

| |
|---|
| 74 % Postcentral gyrus R |
| 55 % Fusiform R/ 41 % Inferior temporal gyrus R |
| 33 % Fusiform R / 27 % Inferior temporal gyrus R |
| 70 % Inferior temporal gyrus R |
| 67 % Middle occipital gyrus R |
| 50 % Hippocampus R / 9 % Parahippocampal gyrus R |
| 46 % Caudate nucleus R / 6 % Thalamus R |
| 84 % Lenticular nucleus, putamen R |
| 41 % Thalamus R / 8 % Hippocampus R |
| 40 % Superior frontal gyrus,medial L/ 30 % Superior frontal gyrus, dorsolateral L |
| 61 % Postcentral gyrus L |
| 42 % Rolandic operculum L/ 39 % Insula L |
| 83 % Inferior temporal gyrus L |
| 63 % Inferior temporal gyrus L |
| 37 % Median cingulate and paracingulate gyri L / 36 % Anterior cingulate and paracingulate gyri L |
| 39 % Median cingulate and paracingulate gyri L/ 22 % Posterior cingulate gyrus L |
| 55 % Posterior cingulate gyrus L / 21 % Precuneus L |
| 42 % Hippocampus L/ 5 % Thalamus L |
| 30 % Hippocampus L/ 21 % Parahippocampal gyrus L |
| 55 % Hippocampus L / 13 % Parahippocampal gyrus L |
| 28 % Caudate nucleus L/ 15 % Olfactory cortex L |

Table S4: **ROIs with the highest absolute difference in the bifurcation parameters when comparing normal wakefulness, W and recovery from anaesthesia R.** All the ROIs showed an absolute difference greater than the threshold given by the sum of the mean and standard deviation of the absolute differences corresponding to all the ROIs. The percentage corresponds to the covered part of the ROI in the Automated Anatomical Labeling (AAL) parcellation code.

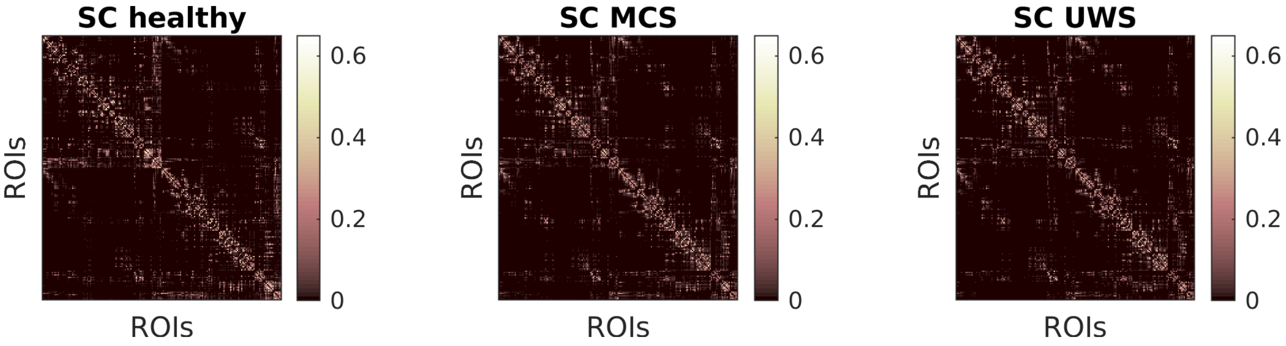


Figure S9: **Examples of structural connectivity (SC) matrices corresponding to the group average of each brain state.** The weights of the SC, $C_{ij}$ correspond to the number of connections between area $i$ with area $j$, corrected to be symmetric.

| | |
|---|---|
| 41 % Thalamus R / 8 % Hippocampus R | 44 % Frontal Medial L / 43 % Superior frontal gyrus, dorsolateral L |
| 55 % Middle temporal gyrus L/ 27 % Angular gyrus L | 50 % Superior frontal gyrus,medial L/ 47 % Anterior cingulate and paracingulate gyri L |
| 80 % Median cingulate and paracingulate gyri R | 42 % Rolandic operculum L / 39 % Insula L |
| 51 % Thalamus L / 2 % Caudate nucleus L | 75 % Precuneus L |
| 42 % Hippocampus L / 5 % Thalamus L | 71 % Fusiform L |
| 66 % Median cingulate and paracingulate gyri L | 57 % Inferior parietal gyrus L / 36 % Supramarginal gyrus L |
| 74 % Supplementary motor area R | 82 % Middle temporal gyrus L |
| 58 % Fusiform L / 36 % Inferior temporal gyrus L | 54 % Postcentral gyrus L/ 28 % Inferior parietal gyrus L |
| 44 % Superior temporal gyrus L / 26 % Rolandic operculum L | 50 % Hippocampus R/ 9 % Parahippocampal gyrus R |
| 81 % Anterior cingulate and paracingulate gyri R | 88 % Lingual gyrus L |
| 52 % Median cingulate and paracingulate gyri L/ 45 % Supplementary motor area L | 60 % Supplementary motor area R/ 22 % Median cingulate and paracingulate gyri R |
| 44 % Frontal Med VMPFC R/ 31 % Anterior cingulate and paracingulate gyri R | 46 % Inferior frontal gyrus, orbital part L / 36 % Insula L |
| 53 % Thalamus L/ 0 % Thalamus R | 55 % Posterior cingulate gyrus L/ 21 % Precuneus L |
| 33 % Anterior cingulate and paracingulate gyri L / 23 % Gyrus rectus L | 52 % Superior frontal gyrus, dorsolateral R/ 41 % Superior frontal gyrus,medial R |
| 57 % Gyrus rectus R / 23 % Frontal Med R | 67 % Calcarine fissure and surrounding cortex L |
| 65 % Inferior temporal gyrus L | 64 % Middle temporal gyrus R |
| 52 % Thalamus R | 62 % Temporal pole: middle temporal gyrus R |
| 46 % Caudate nucleus R/ 6 % Thalamus R | 57 % Precuneus R/ 32 % Calcarine fissure and surrounding cortex R |
| 56 % Median cingulate and paracingulate gyri R/ 26 % Anterior cingulate and paracingulate gyri R | 63 % Inferior temporal gyrus L |
| 80 % Precuneus R | 30 % Hippocampus L/ 21 % Parahippocampal gyrus L |
| 54 % Frontal Med VMPFC L/ 31 % Anterior cingulate and paracingulate gyri L | 42 % Gyrus rectus R/ 19 % Olfactory cortex R |
| 56 % Gyrus rectus L/ 13 % Olfactory cortex L | 62 % Middle temporal gyrus L |
| 49 % Inferior parietal gyrus L/ 38 % Postcentral gyrus L | 49 % Inferior occipital gyrus L/ 33 % Fusiform gyrus L |
| 81 % Median cingulate and paracingulate gyri R | 36 % Thalamus R / 9 % Lingual R |
| 47 % Supramarginal gyrus R/ 35 % Postcentral gyrus R | 87 % Angular gyrus R |
| 52 % Caudate nucleus L/ 2 % Thalamus L | 51 % Lingual gyrus R/ 33 % Fusiform gyrus R |
| 65 % Cuneus R | 50 % Precental gyrus L/ 30 % Inferior frontal gyrus, opercular part L |
| 62 % Caudate nucleus R | 57 % Caudate nucleus R/ 11 % Olfactory cortex R |
| 40 % Fusiform gyrus L/ 31 % Lingual gyrus L | 45 % Precuneus L/ 34 % Calcarine fissure and surrounding cortex L |
| 50 % Anterior cingulate and paracingulate gyri L/ 39 % Superior frontal gyrus, medial L | 46 % Supramarginal gyrus R/ 34 % Inferior parietal gyrus R |
| 59 % Inferior frontal gyrus, orbital part R/ 20 % Insula R | 47 % Postcentral gyrus R/ 23 % Inferior parietal gyrus R |
| 72 % Hippocampus R | 34 % Calcarine fissure and surrounding cortex L/ 26 % Middle occipital gyrus L |
| 46 % Thalamus L/ 1 % Lingual gyrus L | 61 % Inferior temporal gyrus L |

Table S5: **ROIs that show a significant decrease ($p < 0.05$) when comparing the values of the strength obtained from the control subjects and the MCS patients.** Statistics were assessed using Ranksum Wilconsom test, followed by FDR p-value correction. The percentage corresponds to the covered part of the ROI in the Automated Anatomical Labeling (AAL) parcellation code.

| | |
|---|---|
| 41 % Thalamus R/ 8 % Hippocampus R | 47 % Supramarginal gyrus R/ 35 % Postcentral R |
| 51 % Thalamus L/ 2 % Caudate nucleus L | 61 % Inferior temporal gyrus L |
| 53 % Thalamus L/ 0 % Thalamus R | 49 % Inferior occipital gyrus L/ 33 % Fusiform gyrus L |
| 50 % Precental gyrus L/ 30 % Inferior frontal gyrus, opercular part L | 48 % Fusiform gyrus R/ 38 % Inferior temporal gyrus R |
| 80 % Precuneus R | 70 % Superior parietal gyrus R |
| 52 % Thalamus R | 58 % Supplementary motor area R/<br>42 % Superior frontal gyrus, dorsolateral R |
| 42 % Hippocampus L/ 5 % Thalamus L | 65 % Cuneus R |
| 75 % Precuneus L | 36 % Thalamus R/ 9 % Lingual gyrus R |
| 33 % Anterior cingulate and paracingulate gyri L/ 23 % Gyrus rectus L | 66 % Median cingulate and paracingulate gyri L |
| 63 % Postcentral gyrus L 46 % Inferior frontal gyrus, orbital part L | 36 % Insula L |
| 54 % Postcentral gyrus L/ 28 % Inferior parietal gyrus L | 63 % Inferior frontal gyrus, triangular gyrus part L |
| 80 % Median cingulate and paracingulate gyri R | 46 % Inferior temporal gyrus L / 41 % Fusiform gyrus L |
| 46 % Caudate nucleus R/ 6 % Thalamus R | 59 % Fusiform gyrus L/ 21 % Inferior temporal gyrus L |
| 52 % Caudate nucleus L/ 2 % Thalamus L | 45 % Inferior frontal gyrus, triangular part R/<br>28 % Inferior frontal gyrus, orbital part R |
| 44 % Superior frontal gyrus,medial L/<br>43 % Superior frontal gyrus, dorsolateral L | 66 % Inferior temporal gyrus R |
| 56 % Gyrus rectus L/ 13 % Olfactory cortex L | 66 % Middle frontal gyrus L |
| 49 % Inferior parietal gyrus L/ 38 % Postcentral gyrus L | 52 % Superior frontal gyrus, dorsolateral R/<br>41 % Superior frontal gyrus,medial R |
| 62 % Caudate nucleus R | 57 % Gyrus rectus R / 23 % Frontal Med R |
| 62 % Middle temporal gyrus L | 56 % Median cingulate and paracingulate gyri R<br>/ 26 % Anterior cingulate and paracingulate gyri R |
| 81 % Median cingulate and paracingulate gyri R | 53 % Fusiform gyrus R/ 16 % Lingual gyrus R |
| 44 % Medial frontal gyrus R/<br>31 % Anterior cingulate and paracingulate gyri R | 50 % Hippocampus R/ 9 % Parahippocampal gyrus R |
| 55 % Middle temporal gyrus L/ 27 % Angular gyrus L | 52 % Median cingulate and paracingulate gyri L/<br>45 % Supplementary motor area L |
| 72 % Hippocampus R | 66 % Postcentral gyrus R |
| 57 % Precuneus R/ 32 % Calcarine fissure and surrounding cortex R | 47 % Postcentral gyrus R/ 23 % Inferior parietal gyrus R |
| 58 % Fusiform gyrus L/ 36 % Inferior temporal gyrus L | 28 % Caudate nucleus L/ 15 % Olfactory cortex L |
| 40 % Fusiform gyrus L/ 31 % Lingual L | 63 % Postcentral gyrus R |
| 44 % Inferior frontal gyrus, opercular part R/<br>34 % Inferior frontal gyrus, triangular part R | 55 % Posterior cingulate gyrus L/ 21 % Precuneus L |
| 57 % Inferior parietal gyrus L/ 36 % Supramarginal gyrus L | 28 % Amygdala R / 27 % Temporal pole: superior temporal gyrus R |
| 59 % Inferior frontal gyrus, orbital part R/ 20 % Insula R | 62 % Temporal pole: middle temporal gyrus R |
| 46 % Thalamus L/ 1 % Lingual gyrus L | 81 % Supplementary motor area L |
| 42 % Superior occipital gyrus R/ 26 % Cuneus R | 34 % Insula R/ 22 % Lenticular nucleus, putamen R |
| 51 % Lingual gyrus R/ 33 % Fusiform gyrus R | 42 % Rolandic operculum L / 39 % Insula L |
| 57 % Superior frontal gyrus,medial L/<br>41 % Superior frontal gyrus, dorsolateral L | 76 % Calcarine fissure and surrounding cortex R |
| 75 % Middle frontal gyrus L | 51 % Inferior frontal gyrus, orbital part L / 31 % Frontal SomeOrb L |
| 48 % Fusiform gyrus R/ 29 % Inferior temporal gyrus R | 87 % Caudate nucleus L |
| 44 % Superior temporal gyrus L/ 26 % Rolandic operculum L | 50 % Superior frontal gyrus,medial L/<br>47 % Anterior cingulate and paracingulate gyri L |
| 71 % Fusiform gyrus L | 55 % Hippocampus L / 13 % ParaHippocampal L |
| 67 % Calcarine fissure and surrounding cortex L | 65 % Inferior temporal gyrus L |
| 88 % Lingual gyrus L | 84 % Middle frontal gyrus R |
| 19 % Lenticular nucleus, putamen R/ 14 % Caudate nucleus R | 52 % Precental gyrus R/ 22 % Inferior frontal gyrus, opercular part R |
| 54 % Frontal Med VMPFC L/<br>31 % Anterior cingulate and paracingulate gyri L | 78 % Lingual gyrus R |
| 81 % Anterior cingulate and paracingulate gyri R | 73 % Middle occipital gyrus L |
| 30 % Hippocampus L/ 21 % Parahippocampal gyrus L | 74 % Postcentral gyrus R |
| 39 % Fusiform gyrus R/ 25 % Inferior occipital gyrus R | 44 % Calcarine fissure and surrounding cortex R/ 6 % Lingual gyrus R |
| 56 % Temporal pole: middle temporal gyrus R/ 20 % Fusiform gyrus R | 33 % Fusiform gyrus R / 27 % Inferior temporal gyrus R |
| 59 % Angular gyrus L/ 26 % Inferior parietal gyrus L | 82 % Middle temporal gyrus L |
| 67 % Inferior temporal gyrus R | |

Table S6: **ROIs that show a significant decrease ($p < 0.05$) when comparing the values of the strength obtained from the control subjects and UWS patients.** Statistics were assesed using Ranksum Wilconsom test, followed by FDR p-value correction. The percentage corresponds to the covered part of the ROI in the Automated Anatomical Labeling (AAL) parcellation code.

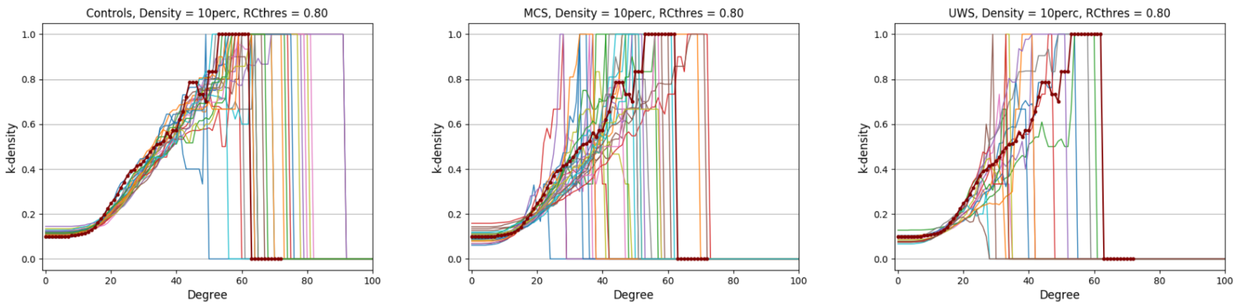


Figure S10: **Rich club extraction based on the k-density for each of the subject**. The curve of the k-density in function of the degree. The ROIs that show a k-density higher than 0.9 where consider to be part of the Rich Club areas. The red line corresponds to the averaged SC of the controls.

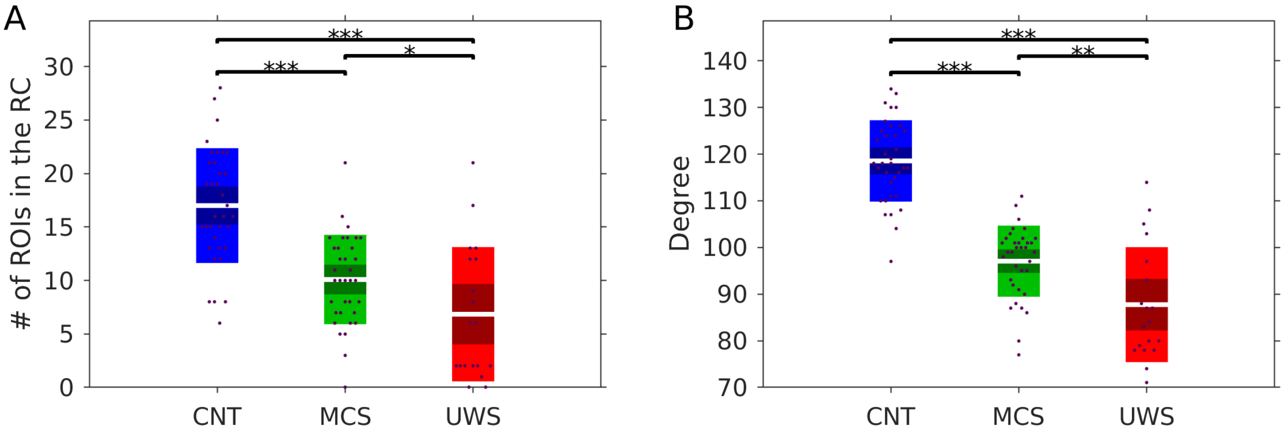


Figure S11: **Structure of the rich club architecture for each subject. A)** Degree distribution of Rich Club ROIs. **B)** Distribution of the number of ROIs considered Rich Club in each group. The number of ROIs changed significantly across groups, decreasing for low level states of consciousness.

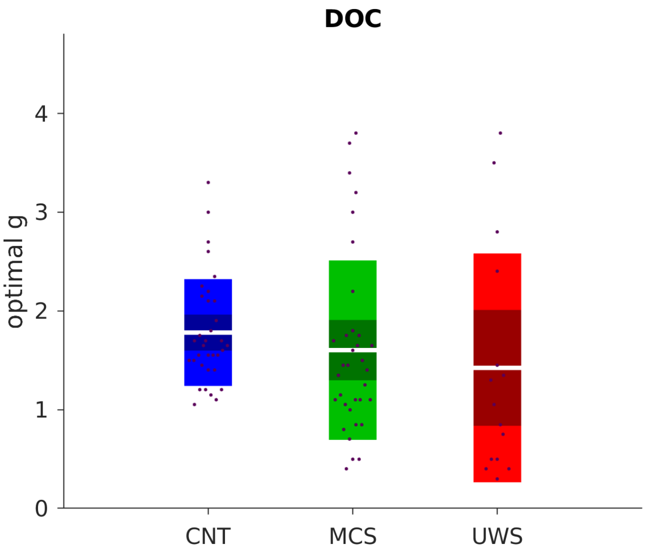


Figure S12: **Whole-brain model global coupling parameter fitting for the individual SC.**Optimal global coupling $g$ for each of the subjects of the DOC dataset using the individual SC to set the interactions between model nodes. One-way-ANOVA p-value: $p_{CNT-MCS} = 0.665$, $p_{CNT-UWS} = 0.446$ and $p_{UWS-MCS} = 0.878$

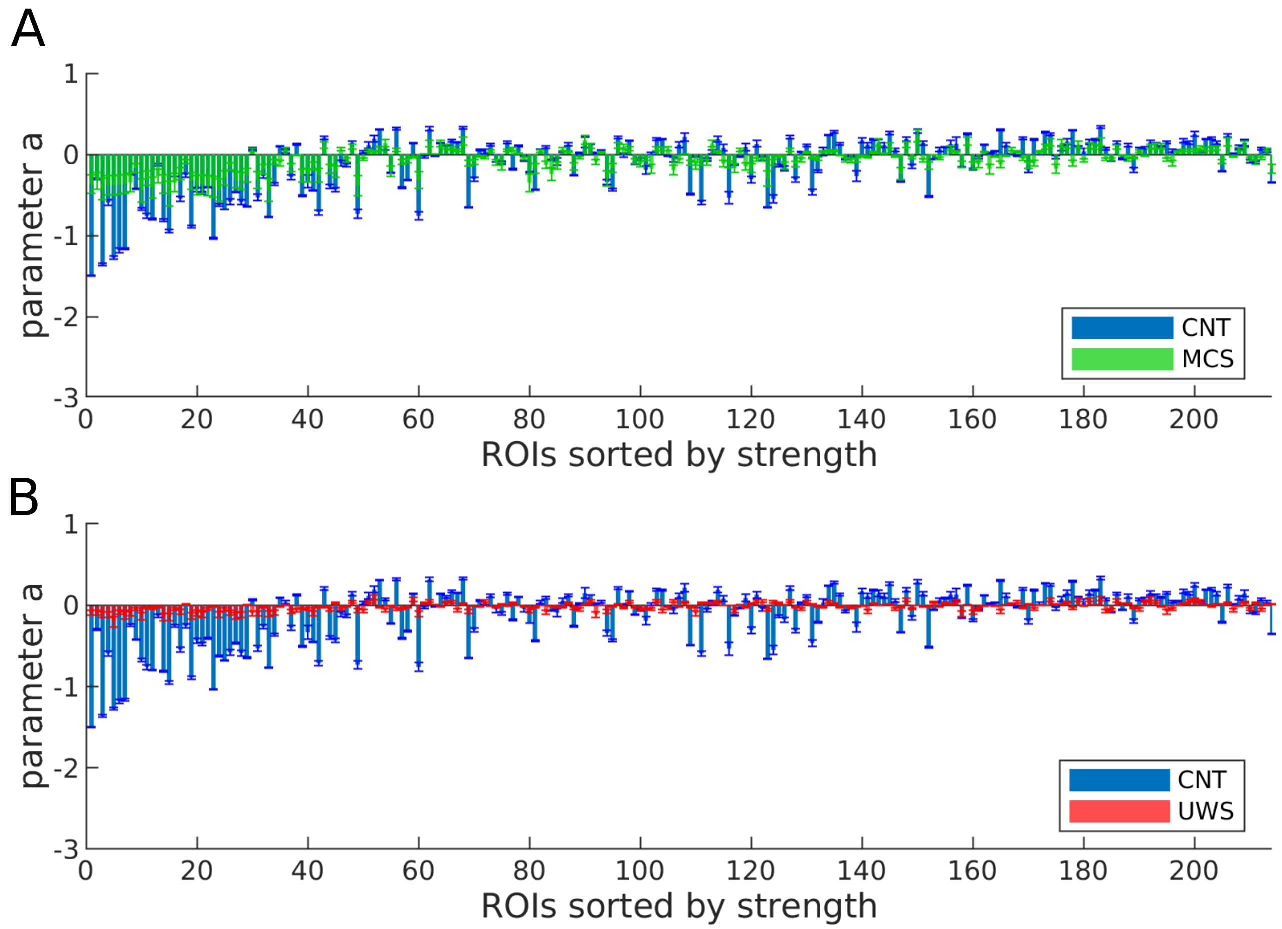


Figure S13: **Local bifurcation parameters of the whole-brain model when using the as SC the patients average SC. A-B)** Bars indicate the mean ± standard error of estimated bifurcation model parameters for each of the 214 nodes (sorted by node strength of the controls SC). Each DOC group results were compared to the wakefulness in heakthy controls.

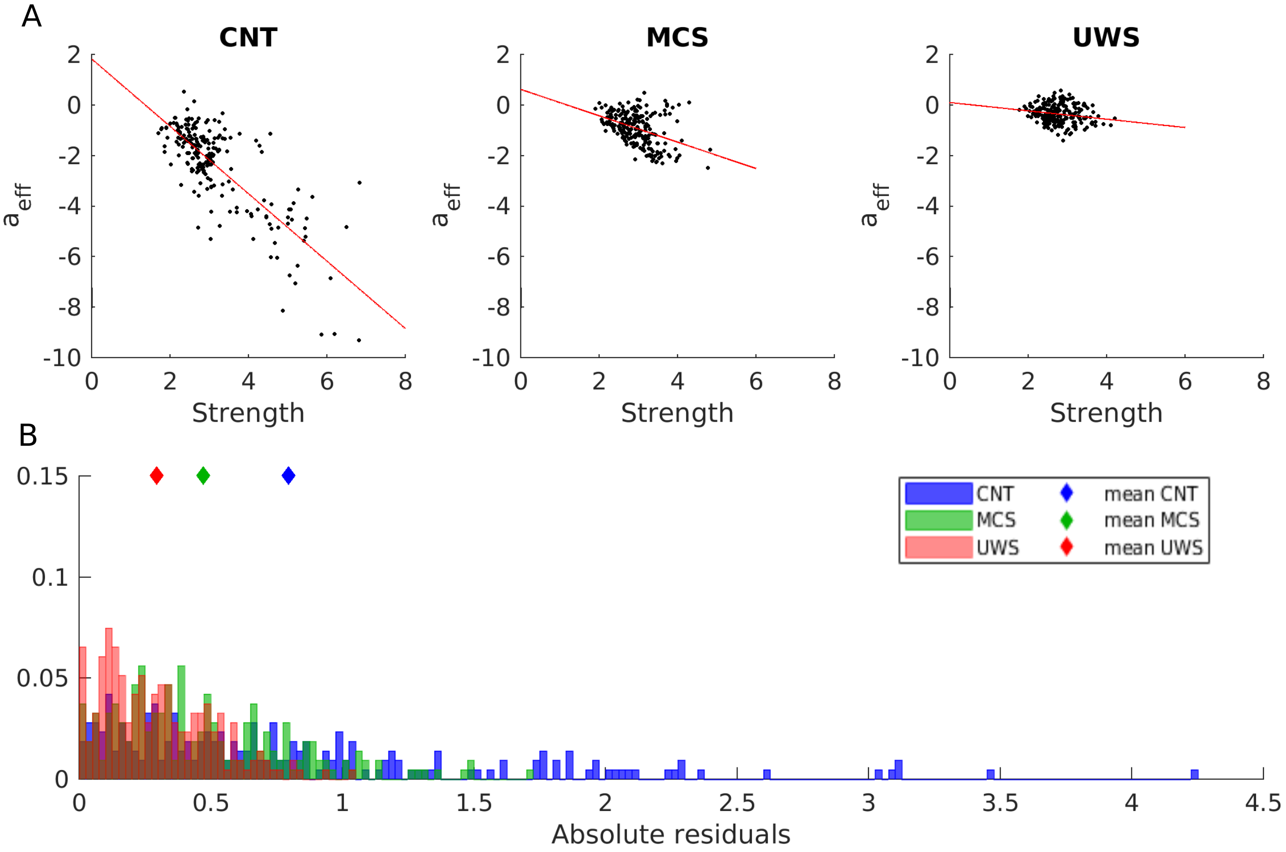


Figure S14: **Relation between the $a_j^{eff}$ and connectivity strength of the SC characterizing each group.** **A)** The effective local bifurcation parameters, $a_j^{eff}$, were estimated using the heterogeneous model and group SC. The $g$ was fixed in all cases using the mean showed in Fig. 2 E. The obtained parameters were compared to the strengths of the nodes $S_j$ in each group SC. The red lines indicate the linear fits. **B)** Distribution of the absolute residuals of each node given by the squared difference between the value of $a_j^{eff}$ and the estimated linear relationship between $a_j^{eff}$ and $S_j$, for each group.

| Condition | Etiology | TSI | Age | Gender | Auditory | Visual | Motor | Verbal | Communication | Arousal | Total CRS-R |
|---|---|---|---|---|---|---|---|---|---|---|---|
| MCS 1 | TBI | 3034 | 34 | F | 3 | 3 | 2 | 2 | 0 | 2 | 12 |
| MCS 2 | TBI | 1294 | 40 | F | 2 | 3 | 2 | 2 | 0 | 2 | 11 |
| MCS 3 | CVA | 13 | 62 | M | 0 | 3 | 2 | 1 | 0 | 1 | 7 |
| MCS 4 | TBI | 589 | 30 | M | 3 | 2 | 2 | 2 | 0 | 1 | 10 |
| MCS 5 | TBI | 28 | 65 | M | 3 | 4 | 3 | 1 | 0 | 2 | 13 |
| MCS 6 | Haemorrhage | 17 | 83 | M | 3 | 0 | 2 | 1 | 0 | 0 | 6 |
| MCS 7 | TBI | 521 | 28 | M | 1 | 3 | 2 | 2 | 0 | 2 | 10 |
| MCS 8 | Epilepsy | 20 | 52 | M | 3 | 3 | 2 | 2 | 1 | 2 | 13 |
| MCS 9 | Haemorrhage | 43 | 67 | M | 2 | 3 | 5 | 2 | 0 | 2 | 14 |
| MCS 10 | TBI | 533 | 47 | M | 3 | 5 | 2 | 1 | 0 | 2 | 13 |
| MCS 11 | CVA | 2639 | 38 | M | 1 | 3 | 2 | 2 | 0 | 1 | 9 |
| MCS 12 | TBI | 2690 | 24 | M | 3 | 3 | 5 | 1 | 0 | 2 | 14 |
| MCS 13 | TBI+anoxia | 401 | 29 | M | 1 | 3 | 2 | 1 | 0 | 2 | 9 |
| MCS 14 | Anoxia | 9900 | 39 | M | 3 | 3 | 5 | 2 | 0 | 2 | 15 |
| MCS 15 | Anoxia | 396 | 57 | M | 3 | 0 | 2 | 2 | 0 | 2 | 9 |
| MCS 16 | TBI+Anoxia | 314 | 26 | F | 3 | 1 | 2 | 1 | 0 | 2 | 9 |
| MCS 17 | TBI | 407 | 31 | M | 0 | 1 | 2 | 2 | 0 | 1 | 6 |
| MCS 18 | Anoxia | 64 | 29 | M | 1 | 3 | 2 | 2 | 0 | 2 | 10 |
| MCS 19 | Haemorrhage | 242 | 46 | F | 2 | 3 | 2 | 1 | 0 | 2 | 10 |
| MCS 20 | Anoxia | 639 | 43 | M | 2 | 3 | 1 | 2 | 0 | 2 | 10 |
| MCS 21 | TBI | 1241 | 53 | M | 0 | 3 | 2 | 2 | 0 | 2 | 9 |
| MCS 22 | TBI | 135 | 51 | M | 3 | 4 | 2 | 1 | 0 | 2 | 12 |
| MCS 23 | TBI | 30 | 67 | F | 0 | 3 | 2 | 0 | 0 | 2 | 7 |
| MCS 24 | Haemorrhage | 1383 | 68 | F | 3 | 1 | 3 | 2 | 0 | 2 | 11 |
| MCS 25 | TBI | 1331 | 35 | M | 3 | 0 | 2 | 1 | 0 | 2 | 8 |
| MCS 26 | TBI - Haemorrhage | 35 | 73 | M | 0 | 2 | 0 | 1 | 0 | 1 | 4 |
| MCS 27 | CVA | 104 | 43 | F | 3 | 1 | 3 | 1 | 0 | 0 | 8 |
| MCS 28 | TBI | 319 | 41 | M | 1 | 0 | 2 | 1 | 0 | 1 | 5 |
| MCS 29 | TBI - Haemorrhage | 255 | 39 | M | 4 | 5 | 4 | 2 | 1 | 2 | 18 |
| MCS 30 | Anoxia | 1482 | 32 | M | 3 | 4 | 2 | 2 | 1 | 2 | 14 |
| MCS 31 | TBI | 641 | 23 | M | 3 | 3 | 0 | 1 | 0 | 2 | 9 |
| MCS 32 | TBI | 37 | 26 | F | 2 | 3 | 3 | 0 | 1 | 1 | 10 |
| MCS 33 | Haemorrhage | 389 | 59 | F | 2 | 1 | 2 | 1 | 0 | 2 | 8 |

Table S7: **MCS patients' demographic and clinical characteristics.** The table includes condition, etiology (traumatic brain injury (TBI) and cerebral vascular accident (CVA)), time science injury (TSI), age, gender (F=female, M=male), Coma Recovery Scale-Revised (CSR-R) auditory, visual, motor, verbal, communication and arousal subscores and total.

| **Condition** | **Etiology** | **TSI** | **Age** | **Gender** | **Auditory** | **Visual** | **Motor** | **Verbal** | **Communication** | **Arousal** | **Total CRS-R** |
|---|---|---|---|---|---|---|---|---|---|---|---|
| UWS 1 | Anoxia | 2890 | 49 | M | 1 | 1 | 1 | 2 | 0 | 2 | 7 |
| UWS 2 | TBI | 283 | 52 | F | 1 | 0 | 2 | 2 | 0 | 1 | 6 |
| UWS 3 | Anoxia | 743 | 30 | M | 1 | 0 | 2 | 1 | 0 | 2 | 6 |
| UWS 4 | Anoxia | 92 | 74 | M | 1 | 0 | 1 | 1 | 0 | 1 | 4 |
| UWS 5 | Haemorrhage | 43 | 64 | M | 1 | 0 | 2 | 1 | 0 | 1 | 5 |
| UWS 6 | Anoxia | 18 | 20 | M | 1 | 0 | 0 | 1 | 0 | 1 | 3 |
| UWS 7 | Anoxia | 1683 | 39 | F | 1 | 0 | 2 | 1 | 0 | 2 | 6 |
| UWS 8 | Anoxia | 38 | 50 | F | 0 | 0 | 0 | 2 | 0 | 1 | 3 |
| UWS 9 | Anoxia | 50 | 69 | F | 0 | 1 | 2 | 1 | 0 | 1 | 5 |
| UWS 10 | Anoxia | 129 | 49 | F | 1 | 0 | 0 | 1 | 0 | 2 | 4 |
| UWS 11 | TBI | 24 | 58 | M | 0 | 1 | 2 | 0 | 0 | 1 | 4 |
| UWS 12 | Anoxia | 335 | 40 | F | 1 | 0 | 2 | 1 | 0 | 2 | 6 |
| UWS 13 | Anoxia | 7814 | 34 | M | 1 | 0 | 1 | 1 | 0 | 2 | 5 |
| UWS 14 | Anoxic Asphyxia | 304 | 60 | M | 1 | 1 | 1 | 1 | 0 | 2 | 6 |
| UWS 15 | Anoxia | 30 | 44 | M | 1 | 1 | 1 | 1 | 0 | 1 | 5 |

Table S8: **UWS patients' demographic and clinical characteristics.** The table includes condition, etiology (traumatic brain injury (TBI)), time science injury (TSI), age, gender (F=female, M=male), Coma Recovery Scale-Revised (CSR-R) auditory, visual, motor, verbal, communication and arousal subscores and total.

| ROI | Corresponding label in AAL atlas | ROI | Corresponding label in AAL atlas |
|---|---|---|---|
| 1 | 38 % Superior frontal gyrus, orbital part [6] / 25 % Middle frontal gyrus, orbital part [10] | 55 | 67 % Inferior temporal gyrus [90] |
| 2 | 42 % Gyrus rectus [28] / 19 % Olfactory cortex [22] | 56 | 69 % Inferior temporal gyrus [90] |
| 3 | 57 % Gyrus rectus [28] / 23 % Superior frontal gyrus, medial orbital [26] | 57 | 67 % Inferior temporal gyrus [90] |
| 4 | 69 % Superior frontal gyrus, orbital part [6] | 58 | 48 % Fusiform gyrus [56] / 38 % Inferior temporal gyrus [90] |
| 5 | 44 % Superior frontal gyrus, medial orbital [26] / 31 % Anterior cingulate and paracingulate gyri [32] | 59 | 55 % Fusiform gyrus [56] / 41 % Inferior temporal gyrus [90] |
| 6 | 34 % Superior frontal gyrus, dorsolateral [4] / 33 % Superior frontal gyrus, medial [24] | 60 | 33 % Fusiform gyrus [56] / 27 % Inferior temporal gyrus [90] |
| 7 | 41 % Middle frontal gyrus, orbital part [10] / 29 % Middle frontal gyrus [8] | 61 | 50 % Superior temporal gyrus [82] / 28 % Rolandic operculum [18] |
| 8 | 38 % Inferior frontal gyrus, orbital part [16] / 26 % Middle frontal gyrus, orbital par [10] | 62 | 28 % Superior temporal gyrus [82] / 25 % Rolandic operculum [18] |
| 9 | 61 % Middle frontal gyrus [8] | 63 | 52 % Superior temporal gyrus [82] / 48 % Middle temporal gyrus [86] |
| 10 | 60 % Superior frontal gyrus, medial[24] / 20 % Anterior cingulate and paracingulate gyri [32] | 64 | 66 % Middle temporal gyrus [86] |
| 11 | 84 % Middle frontal gyrus [8] | 65 | 64 % Middle temporal gyrus [86] |
| 12 | 52 % Superior frontal gyrus, dorsolateral [4] / 41 % Superior frontal gyrus, medial [24] | 66 | 66 % Inferior temporal gyrus [90] |
| 13 | 51 % Middle frontal gyrus [8] / 38 % Superior frontal gyrus, dorsolateral [4] | 67 | 39 % Fusiform gyrus [56] / 25 % Inferior occipital gyrus [54] |
| 14 | 85 % Middle frontal gyrus [8] | 68 | 53 % Fusiform gyrus [56] / 16 % Lingual gyrus [48] |
| 15 | 56 % Median cingulate and paracingulate gyri [34] / 26 % Anterior cingulate and paracingulate gyri [32] | 69 | 51 % Inferior temporal gyrus [90] / 47 % Middle temporal gyrus [86] |
| 16 | 45 % Inferior frontal gyrus, triangular part [14] / 28 % Inferior frontal gyrus, orbital part [16] | 70 | 70 % Inferior temporal gyrus [90] |
| 17 | 56 % Inferior frontal gyrus, orbital part [16] / 29 % Middle frontal gyrus, orbital part [10] | 71 | 48 % Fusiform gyrus [56] / 29 %Inferior temporal gyrus [90] |
| 18 | 59 %Inferior frontal gyrus, orbital part [16] / 20 % Insula [30] | 72 | 51 % Lingual gyrus [48] / 33 % Fusiform gyrus [56] |
| 19 | 67 % Inferior frontal gyrus, triangular part [14] | 73 | 67 % Middle occipital gyrus [52] |
| 20 | 57 % Insula [30] / 21 % Inferior frontal gyrus, triangular part [14] | 74 | 34 % Middle occipital gyrus [52] / 32 % Middle temporal gyrus [86] |
| 21 | 45 % Inferior frontal gyrus, opercular part [12] / 34 % Precental gyrus [2] | 75 | 42 % Superior occipital gyrus [50] / 26 % Cuneus [46] |
| 22 | 44 % Inferior frontal gyrus, opercular part [12] / 34 % Inferior frontal gyrus, triangular part [14] | 76 | 48 % Lingual gyrus [48] / 17 % Fusiform gyrus [56] |
| 23 | 74 % Postcentral gyrus [58] | 77 | 65 % Cuneus [46] |
| 24 | 57 % Supplementary motor area [20] / 39 % Paracentral Lobule [70] | 78 | 37 %Middle occipital gyrus [52] / 20 % Superior occipital gyrus [50] |
| 25 | 74 % Supplementary motor area [20] | 79 | 78 % Lingual gyrus [48] |
| 26 | 51 % Precental gyrus [2] / 43 % Superior frontal gyrus, dorsolateral [4] | 80 | 39 % Calcarine fissure and surrounding cortex [44] / 27 % Cuneus [46] |
| 27 | 81 % Precental gyrus [2] | 81 | 59 %Inferior occipital gyrus [54] / 23 % Lingual gyrus [48] |
| 28 | 60 % Supplementary motor area [20] / 22 % Median cingulate and paracingulate gyri [34] | 82 | 76 % Calcarine fissure and surrounding cortex [44] |
| 29 | 58 % Supplementary motor area [20] / 42 % Superior frontal gyrus, dorsolateral [4] | 83 | 81 % Anterior cingulate and paracingulate gyri [32] |
| 30 | 47 % Middle frontal gyrus [8] / 40 % Superior frontal gyrus, dorsolateral [4] | 84 | 81 % Median cingulate and paracingulate gyri [34] |
| 31 | 52 % Precental gyrus [2] / 22 % Inferior frontal gyrus, opercular part [12] | 85 | 44 % Posterior cingulate gyrus [36] / 39 % Median cingulate and paracingulate gyri [34] |
| 32 | 45 % Precental gyrus [2] / 25 % Middle frontal gyrus [8] | 86 | 57 % Precuneus [68] / 32 % Calcarine fissure and surrounding cortex [44] |
| 33 | 66 % Postcentral gyrus [58] | 87 | 44 % Calcarine fissure and surrounding cortex [44] / 6 % Lingual gyrus [48] |
| 34 | 66 % Insula [30] | 88 | 42 % Median cingulate and paracingulate gyri [34] |
| 35 | 56 % Insula [30] / 24 % Rolandic operculum [18] | 89 | 80 % Median cingulate and paracingulate gyri [34] |
| 36 | 52 % Insula [30] / 36 %Inferior frontal gyrus, orbital part [16] | 90 | 79 % Precuneus [68] |
| 37 | 34 % Insula [30] / 22 % Lenticular nucleus, putamen [74] | 91 | 46 % Precuneus [68] / 37 % Median cingulate and paracingulate gyri [34] |
| 38 | 47 % Postcentral gyrus [58] / 23 % Inferior parietal, but supramarginal and angular gyri [62] | 92 | 28 % Amygdala [42] / 27 % Temporal pole: superior temporal gyrus [84] |
| 39 | 63 % Postcentral gyrus [58] | 93 | 50 % Hippocampus [38] / 9 % ParaHippocampal gyrus [40] |
| 40 | 57 % Rolandic operculum [18] / 34 % Insula [30] | 94 | 72 % Hippocampus [38] |
| 41 | 70 % Superior parietal gyrus [60] | 95 | 36 % ParaHippocampal gyrus [40] / 36 % Hippocampus [38] |
| 42 | 49 % Precuneus [68] / 40 % Cuneus [46] | 96 | 60 % ParaHippocampal gyrus [40] / 28 % Fusiform gyrus [56] |
| 43 | 37 % Superior parietal gyrus [60] / 23 % Inferior parietal, but supramarginal and angular gyri [62] | 97 | 67 % ParaHippocampal gyrus [40] |
| 44 | 80 % Precuneus [68] | 98 | 49 % Lingual gyrus [48] / 23 % Precuneus [68] |
| 45 | 47 % Supramarginal gyrus [64] / 35 % Postcentral gyrus [58] | 99 | 30 % Hippocampus [38] / 15 % Amygdala [42] |
| 46 | 53 % Superior temporal gyrus [82] / 37 % SupraMarginal [64] | 100 | 62 % Caudate nucleus [72] |
| 47 | 46 % Supramarginal gyrus [64] / 34 % Inferior parietal, but supramarginal and angular gyri [62] | 101 | 46 % Caudate nucleus [72] / 6 % Thalamus [78] |
| 48 | 87 % Angular gyrus [66] | 102 | 57 % Caudate nucleus [72] / 11 % Olfactory cortex [22] |
| 49 | 73 % Middle occipital gyrus [52] | 103 | 84 % Lenticular nucleus, putamen [74] |
| 50 | 87 % Middle temporal gyrus [86] | 104 | 19 % Lenticular nucleus, putamen [74] / 14 % Caudate nucleus [72] |
| 51 | 56 % Temporal pole: middle temporal gyrus [88] / 20 % Fusiform gyrus [56] | 105 | 36 % Thalamus [78] / 9 % Lingual gyrus [48] |
| 52 | 62 % Temporal pole: middle temporal gyrus [88] | 106 | 41 % Thalamus [78] / 8 % Hippocampus [38] |
| 53 | 45 % Temporal pole: superior temporal gyrus [84] / 34 % Temporal pole: middle temporal gyrus [88] | 107 | 52 % Thalamus [78] |
| 54 | 58 % Middle temporal gyrus [86] / 32 % Superior temporal gyrus [82] | | |

Table S9: **Correspondences of the Shen parcellation ROIs to the AAL atlas.**ROIs in the Shen atlas for one hemisphere (same parcellation for both hemispheres) and each correspondence to the AAL atlas. The percentage corresponds to the covered part of the ROI in the Automated Anatomical Labeling (AAL) parcellation code.
